# Genome assembly of the A-group *Wolbachia* in *Nasonia oneida* and phylogenomic analysis of *Wolbachia* strains reveals patterns of genome evolution and lateral gene transfer

**DOI:** 10.1101/508408

**Authors:** Xiaozhu Wang, Xiao Xiong, Wenqi Cao, Chao Zhang, John H. Werren, Xu Wang

## Abstract

*Wolbachia* are obligate intracellular bacteria which commonly infect various nematode and arthropod species. Based on depth differences, we assembled the genome of *Wolbachia* in the parasitoid jewel wasp species *Nasonia oneida* (*w*One), using 10X Genomics Chromium linked-read technology. The final draft assembly consists of 1,293,406 bp in 47 scaffolds with 1,114 coding genes and 97.01% genome completeness assessed by checkM. *w*One is the A1 strain previously reported in *N. oneida*, and pyrosequencing confirms that the wasp strain lacks A2 and B types, which were likely lost during laboratory culturing. Polymorphisms identified in the *w*OneA1 genome have elevated read depths, indicating recent gene duplications rather that strain variation. These polymorphisms are enriched in nonsynonymous changes in 27 coding genes, including phase baseplate assembly proteins and transporter activity related genes. *w*OneA1 is more closely grouped with A-*Wolbachia* in the *Drosophila simulans* (*w*Ha) than A-*Wolbachia* found in wasps. Genome variation was next evaluated in 34 published *Wolbachia* genomes for 211 single ortholog genes, and revealed six supergroup discordant trees, indicating recombination events not only between A and B supergroups, but also between A and E supergroups. Comparisons of strain divergence using the five genes of the Multi Locus Strain Typing (MLST) system show a high correlation (rho=0.98) between MLST and whole genome divergences, indicating that MLST is a reliable method for identifying related strains when whole genome data are not available. Assembling bacterial genomes from host genome projects can provide an effective method for sequencing *Wolbachia* genomes and characterizing their diversity.

**Author Summary:** More than half of the arthropod species are infected by the obligated intracellular bacteria *Wolbachia*. As one of the most widespread parasitic microbes, *Wolbachia* mediate important biological processes such as cytoplasmic incompatibility and lateral gene transfer in insects. Their evolutionary relationship has been characterize using five protein-coding and 16S rRNA genes. In this work, we identified 211 conserved single copies genes in 34 genome sequenced *Wolbachia* strains, and we discovered that they maintain the supergroup relationship classified previously based on selected genes. We constructed phylogenetic trees for individual genes and found only six genes display discordant tree structure between supergroups, due to lateral gene transfer and homologous recombination events. But these events are not common (3%) in *Wolbachia* genomes, at least in these conserved single copy genes. In addition to known lateral gene transfer events between A and B supergroups, we identified transfers between A and E supergroups for the first time. Selective maintenance of such transfers suggests possible roles in *Wolbachia* infection related functions. We also found enriched nonsynonymous polymorphisms in *Nasonia oneida Wobachia* genome, and their differences are more likely to result from gene duplications within the strain, rather than strain variation within the parasitoid.

## Background

*Wolbachia*, alphaproteobacterial endosymbionts, are widespread and common in arthropods and filarial nematodes, either as reproductive parasites or mutualists (Fenn and Blaxter 2006; Werren 1997; Werren, et al. 2008). It is estimated that half or more of arthropods are infected with *Wolbachia* (Hilgenboecker, et al. 2008; Zug and Hammerstein 2012), possibly representing a dynamic equilibrium between gain and loss on a global scale (Bailly-Bechet, et al. 2017; Klopfstein, et al. 2018; Werren and Windsor 2000). The widespread distribution of *Wolbachia* is due to horizontal movement of the bacteria between arthropod species, although the routine mode of transmission of these bacteria is vertical through the egg cytoplasm. *Wolbachia* have been found to move across species boundaries through horizontal transfer and by hybrid introgression (Raychoudhury, et al. 2009). The *Wolbachia*-host interaction generally spans a range from reproductive parasitism to mutualism. *Wolbachia* can alter the host reproduction to enhance their own transmission in different ways, such as feminization of genetic males, male-killing, parthenogenetic induction, and cytoplasmic incompatibility (Stouthamer, et al. 1999; Werren, et al. 2008). *Wolbachia pipientis* have been divided into eight supergroups (A-H) based on 16S ribosomal RNA sequences and other sequence information, including six supergroups (A,B and E-H) primarily identified in arthropods and two supergroups (C and D) commonly found in filarial nematodes (Werren, et al. 2008). Supergroup G is now considered as a recombinant of supergroups A and B (Baldo and Werren 2007). A multi-locus strain typing (MLST) system based on five house-keeping genes, (*coxA*, *gatB*, *hcpA*, *ftsZ* and *fbpA*) has been developed for *Wolbachia* (Baldo, et al. 2006b), and is widely used for strain typing and to characterize strain variation within *Wolbachia*. However, the increasing number of genome sequences for *Wolbachia* allows for more detailed characterization of their diversity, including inter-strain recombination events.

The jewel wasp genus of *Nasonia* has been an excellent model for *Wolbachia* research (Bordenstein, et al. 2001; Bordenstein, et al. 2003; Breeuwer and Werren 1993; Perrot-Minnot, et al. 1996; Raychoudhury, et al. 2009). Eleven *Wolbachia* have so far been identified in the four species of *Nasonia*, including two (*w*VitA and *w*VitB) in *N. vitripennis* (NV), three (*w*NgirA1, *w*NgirA2 and *w*NgirB) in *N. giraulti* (NG), three (*w*OneA1, *w*OneA2 and *w*OneB) in *N. oneida* (NO), and three (*w*NlonA, *w*NlonBl and *w*NlonB2) in *N. longicornis* (NL) (Raychoudhury, et al. 2009). These are often maintained as multiple infections within individual wasps of each species. Although these strains belong to two major supergroups (A and B), the *Wolbachia* of each supergroup are not monophyletic, but rather have diverse evolutionary origins, indicating horizontal transfers from divergent host species (Raychoudhury, et al. 2009). The exception to this is the B *Wolbachia* found in *N. longicornis*, *N. giraulti* and *N. oneida*, which are closely related and derived from a common ancestor many of the *Wolbachia* are not monophyletic (which would suggest cospeciation with their hosts).

Genomic studies of *Wolbachia* blossomed in the recent years since the first complete genome of the A-*Wolbachia* parasite of *Drosophila melanogaster* (*w*Mel) published in 2004 (Wu, et al. 2004), followed by the complete genome of D-*Wolbachia* (*w*Bm) in nematode *Brugia malayi* in 2005 (Foster, et al. 2005). A list of sequenced whole genomes of *Wolbachia* is summarized in Table S1. *Wolbachia* genomes are small with a range between 0.9-1.7 Mb. In general, most of the nematode-associated *Wolbachia* have smaller genomes but retain intact metabolic pathways and immunology pathways, which contribute to the mutualistic relationship with the hosts (Darby, et al. 2012; Foster, et al. 2005; Wu, et al. 2004). However, arthropod-associated *Wolbachia* contain more prophage and ankyrin repeat encoding (ANK) gene components, which may reflect their more frequent parasitic lifestyle (Klasson, et al. 2008; Pan, et al. 2008). Furthermore, many studies have claimed lateral gene transfer (LGT) across strains and supergroups, which may be mediated by bacteriophage and lead to the mosaic genomes of *Wolbachia* (Duplouy, et al. 2013; Kent, et al. 2011; Klasson, et al. 2009). Although co-infection of different strains and LGT exist in the same arthropod host, the supergroups may remain genetically distinct clades (Ellegaard, et al. 2013). Another key feature of most *Wolbachia* genomes is the abundance of mobile and repetitive elements, which are different from most *Rickettsiales* (Werren, et al. 2008). Noticeably, the proportion of repetitive elements in genome vary widely among *Wolbachia* strains. For example, 22% of the *w*Ri genome is comprised of repetitive sequences (Klasson, et al. 2009), compared to only 5% in the *w*Bm genome (Foster, et al. 2005). In the jewel wasp (*Nasonia*) species, only two *Wolbachia* strains have been sequenced (*w*VitA (Newton, et al. 2016) and *w*VitB (Kent, et al. 2011)), both from NV. Genome sequence of additional *Wolbachia* strains in the *Nasonia* species complex will facilitate the comparative genomic and evolutionary analyses of this model system.

Because of its endosymbiotic nature, multiple different *Wolbachia* strains can be present in the same host cells, allowing the potential for homologous recombination between strains (Jiggins 2002; Jiggins, et al. 2001). Recombination events in *Wolbachia* have been discovered in *Wsp* (Werren and Bartos 2001) and other genes in Crustaceans (Verne, et al. 2007), mites (Ros, et al. 2012) and insect species including wasps (Baldo, et al. 2006a; Baldo, et al. 2005; Werren and Bartos 2001), ants (Reuter and Keller 2003) and butterflies (Ilinsky and Kosterin 2017), and some of this recombination appears to be phage mediated (Lindsey, et al. 2018). There is no evidence of inter-strain recombination in filarial nematode *Wolbachia* strains (Foster, et al. 2011). Most of the previous research on recombination has focused on five MLST genes, *Wollbachia* surface protein (*wsp*), and 16S, or for a few genomes from the A and B supergroups. Therefore, whole-genome analyses in a large number of *Wolbachia* strains of all supergroups are needed to identify additional homologous recombination and LGT events among *Wolbachia* strains. In this study, we assembled the *Wolbachia* strain in *N. oneida*, performed phylogenomic analyses on 34 genome sequenced *Wolbachia* strains, and analyzed the individual gene tree to identify potential recombination events at the genome level. Relatively low frequencies of intergroup gene transfers were found (6 discordant trees among 211 core single copy genes examined), indicating a general genetic cohesiveness for the A and B supergroups.

## Results

### Assembly of *Wolbachia* genome in the 10X Genomics Chromium sequencing of *N. oneida*

This *Wolbachia* project emerged from an original effort to sequence the genome of the parasitoid wasp *N. oneida* (Raychoudhury, et al. 2010). The *de novo* assembly of the parasitoid wasp NO genome was performed using linked reads generated by 10X Genomics Chromium technology with Supernova 2.1.1 assembler (Weisenfeld, et al. 2017). *Wolbachia* scaffolds were identified and separated from the NO genome assembly using a custom bioinformatics pipeline (Figure 1 and 2A). NO scaffolds were BLATed against bacterial genome database (Kent 2002), and we identified *w*One scaffolds based on the median coverage, GC-content and sequence identity to known *Wolbachia* sequences (Figure 2; see Methods). *w*One scaffolds have an median genome coverage of 59.38X, which is significantly lower compared to 713.59X for the NO genomic scaffolds (*P*-value < 2.2 × 10^−16^, Figure 2B). The mitochondrial scaffold coverage is over 20,000X (Figure 2B). In addition, there is a significant shift in GC-content for the *Wolbachia* genome (35.44%) compared to the host genome average (38.07%, *P*-value = 2.4 × 10^−9^, Figure 2C). The sufficient differences in coverage and GC-content among host genome, host mitochondrial genome and the *Wolbachia* genome allow clear separation of the *w*One scaffolds.

**Figure 1.**
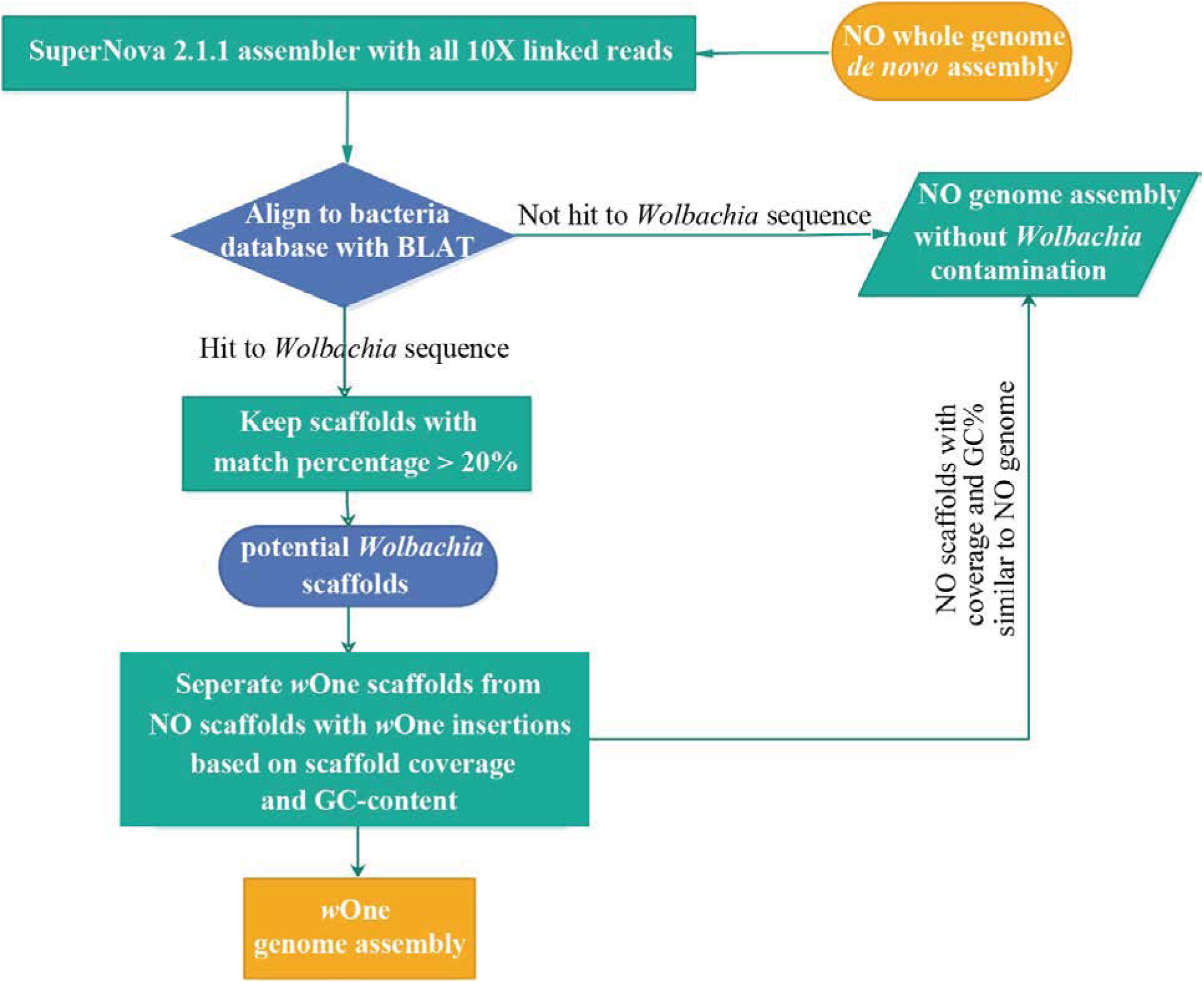
The workflow of *w*One genome assembly using 10X Genomics linked reads.

**Figure 2.**
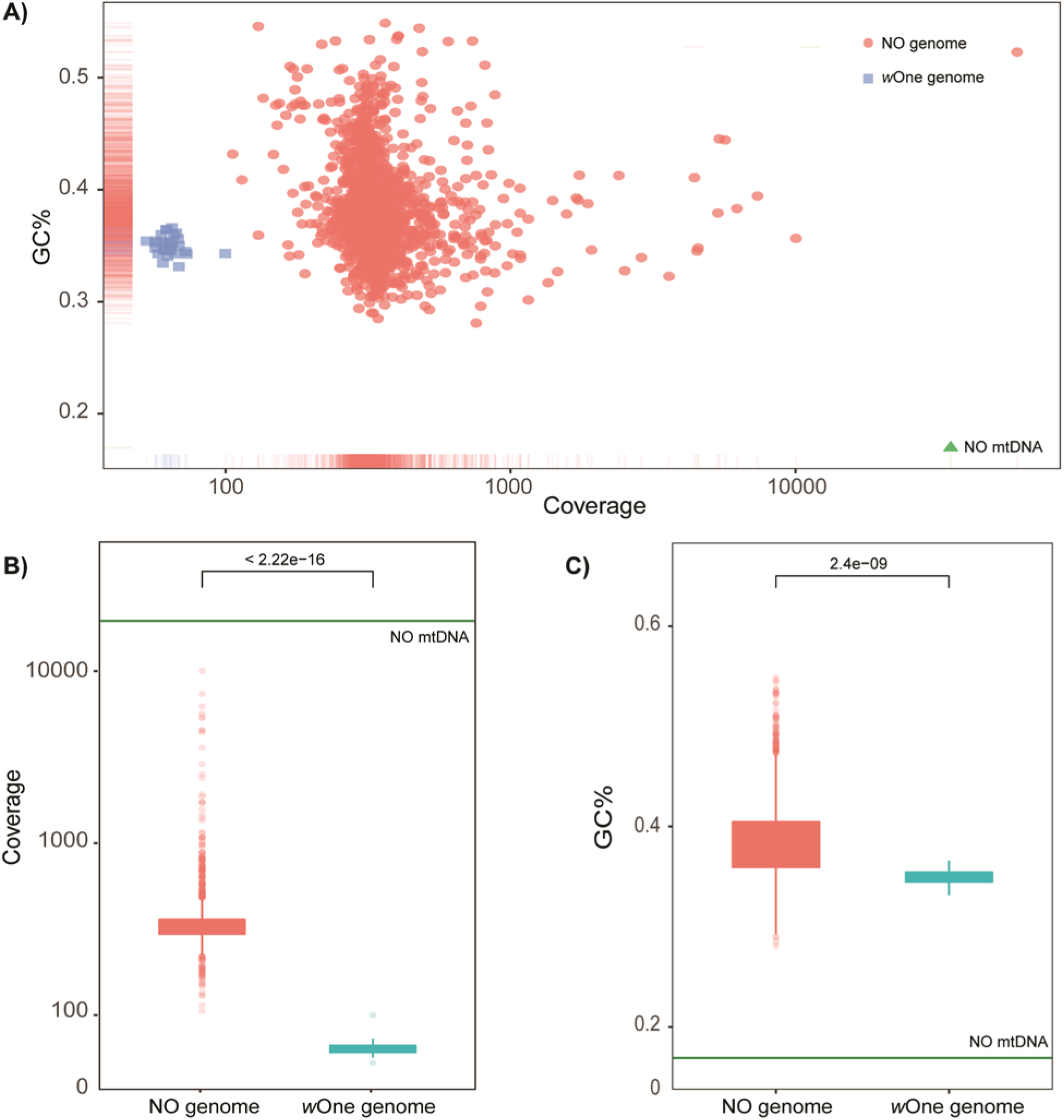
Comparison of median coverage and GC content between *Nasonia oneida* (NO) genome and *w*One genome. (A) scatter plot of median coverage and GC% indicating clear separation of NO and *w*One genomes, blue square represents *w*One genome and red dot represents NO genome, the NO mtDNA was labeled as green triangle; (B) box plot of median coverage between NO and *w*One genomes, green line indicates the coverage of NO mtDNA; (C) box plot of GC% between NO and *w*One genomes, green line indicates the GC% of NO mtDNA.

The *w*One draft genome consists of 1,293,406 nucleotides with 35.44% GC-content (Table 1). This assembly has a total of 47 scaffolds ranging in length from 1,108 to 241,132 bps with a scaffold N50 of 128.97 Kb. A total of 1,114 proteins were annotated in the *w*One genome including rRNAs 5S, 16S and 23S and tRNAs. The number of contigs and scaffolds are fewer than *w*VitA and *w*VitB, and the contig and scaffold N50s are longer (Table 1). The genome completeness is 97.01% accessed by and checkM, which is comparable with *w*VitA and *w*VitB, suggesting high assembly quality. *w*VitA and *w*VitB have slightly higher completeness, but at a cost of 1-2% of contamination (Table 1). The BUSCO completeness is 86.5%, which is typical for complete *Wolbachia* genomes (Sinha, et al. 2019). The 80% of coding genes from *w*VitA, the closest sequenced *Wolbachia* strain in *Nasonia*, were present in *w*One assembly (Figure 3D).

**Figure 3.**
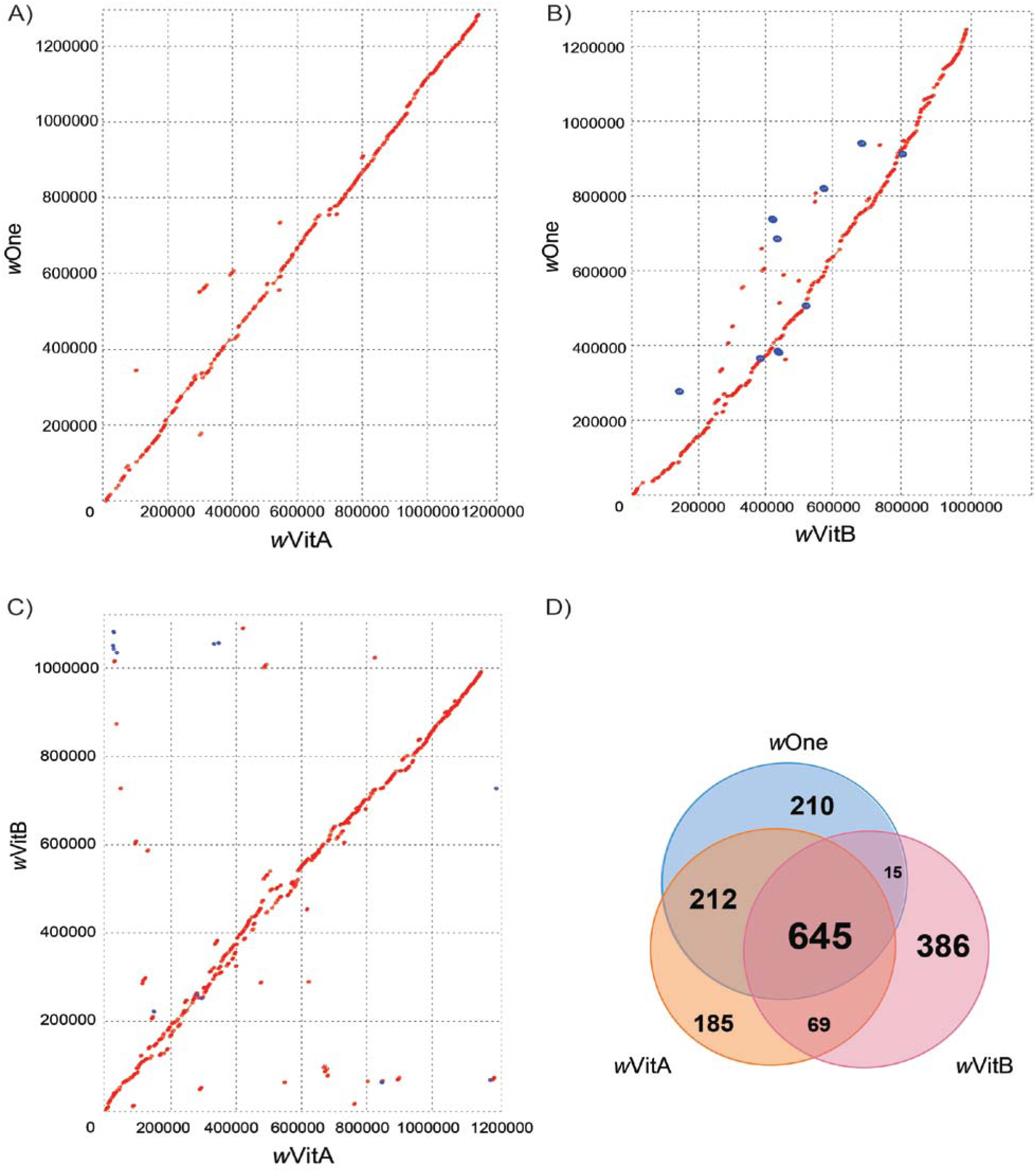
Comparative genomic analysis of *w*One, *w*VitA and *w*VitB genomes. (A) Dot plot showing comparison between *w*One and *w*VitA genomes, red for a forward match and blue for a reverse match; (B) Dot plot showing comparison between *w*One and *w*VitB genomes; (C) Dot plot showing comparison between *w*VitA and *w*VitB genomes; (D) Venn Diagram showing comparison of genes and pseudogenes in *w*One, *w*VitA and *w*VitB.

**Table 1.**
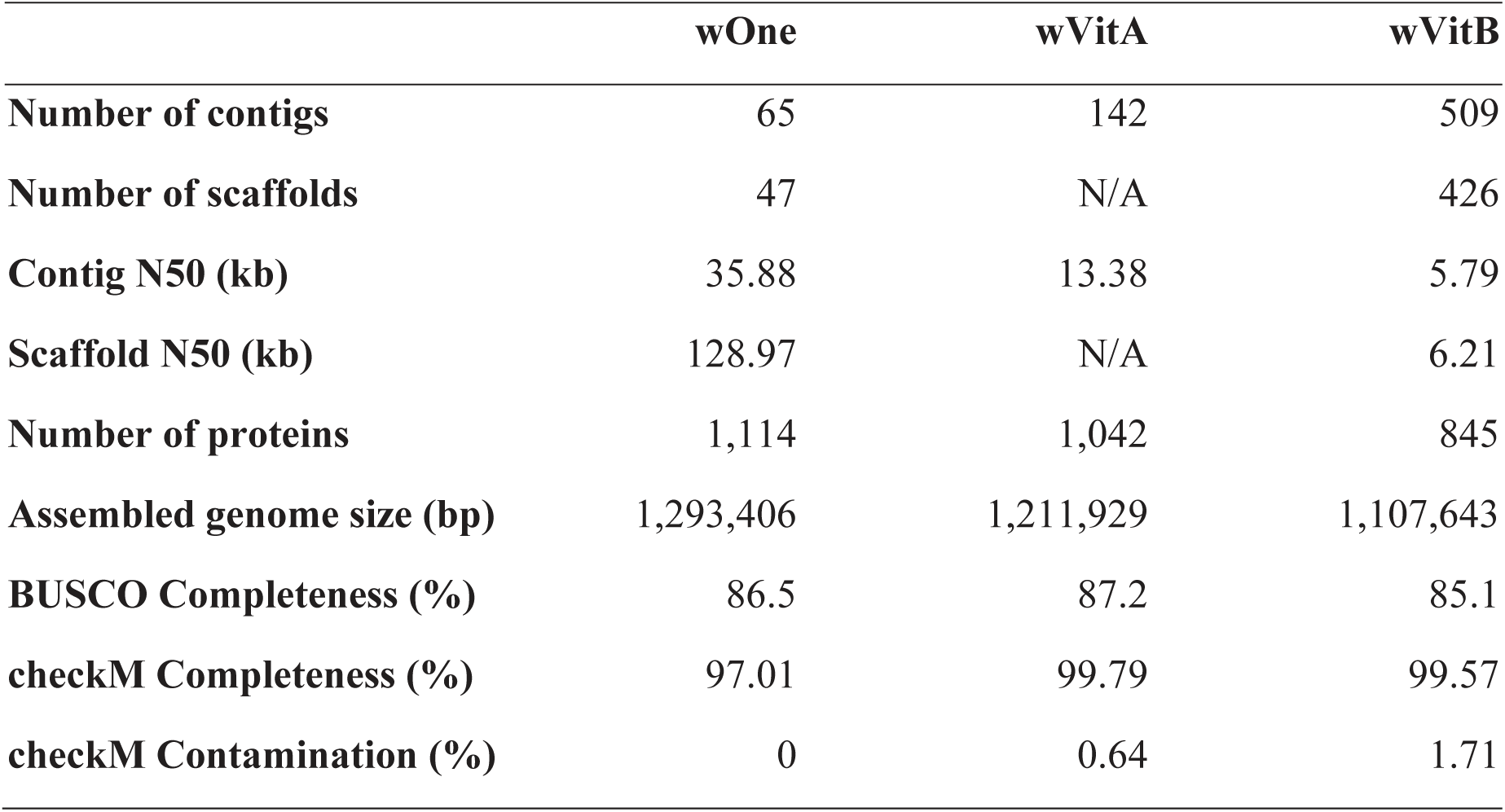
*w*One assembly summary statistics and comparison with *w*VitA and *w*VitB genomes.

|  | ***w*One** | ***w*VitA** | ***w*VitB** |
| --- | --- | --- | --- |
| **Number of contigs** | 65 | 142 | 509 |
| **Number of scaffolds** | 47 | N/A | 426 |
| **Contig N50 (kb)** | 35.88 | 13.38 | 5.79 |
| **Scaffold N50 (kb)** | 128.97 | N/A | 6.21 |
| **Number of proteins** | 1,114 | 1,042 | 845 |
| **Assembled genome size (bp)** | 1,293,406 | 1,211,929 | 1,107,643 |
| **BUSCO Completeness (%)** | 86.5 | 87.2 | 85.1 |
| **checkM Completeness (%)** | 97.01 | 99.79 | 99.57 |
| **checkM Contamination (%)** | 0 | 0.64 | 1.71 |

### Comparative genomic analysis of *Wolbachia* strains in *Nasonia* species

Comparisons of the five MLST genes revealed that the sequenced *Wolbachia* genome is from the strain wOneA1 (Raychoudhury et al 2008). Pairwise alignments were performed to two *Wolbachia* genomes isolated from *N. vitripennis* (*w*VitA and *w*VitB; see Materials and Methods). The assembled genome size for *w*OneA1 is 14% larger than *w*VitB and 6% larger than *w*VitA. The total number of proteins is similar to the *w*VitA and *w*VitB genomes (Table 1). A dot plot revealed a better colinear relationship between *w*OneA1 and *w*VitA, with only a small number of rearrangements around the origin (Figure 3A). A total of 992,405 bps of *w*One genome were aligned *w*VitA genome (covering 81.89%) with an average identity of 96.16%, including 681,482 bps matched in the same orientation and 310,923 bps matched in the reverse orientation. The top 5 longest scaffolds in the *w*One genome (SCAFFOLD1, 2, 3, 4, 5) were aligned to 54.30% *w*VitA genome with an average identity of 96.93%, indicating the high contiguity of wOne genome assembly. However, when comparing *w*OneA1 and *w*VitA genomes with *w*VitB genome respectively, significantly more genome rearrangements and inversions were observed (Figure 3B and C). This is not surprising as *w*VitB belongs to a different supergroup. A total of 671,104 bps of *w*OneA1 genome were aligned *w*VitB genome, including 348,372 bps matched in the same orientation and 322,732 bps matched in the reverse orientation.

When comparing the gene contents of these *Wolbachia* strains, a total of 645 genes were shared among genomes of *w*One, *w*VitA and *w*VitB; 212 more genes were shared between the *w*OneA1 and *w*VitA genomes but not with *w*VitB genome (Figure 3D). Among the 210 *w*One-specific genes, a large fraction belongs to hypothetical protein (N=173) and transposon-related (N=22) genes. Regarding to insertion element (IS), the *w*OneA1 genome contains similar numbers of IS elements when compared to the genomes of *w*VitA and *w*VitB (Table 2). Although *w*VitA and *w*VitB infect the same host NV and *w*OneA1 infect a different host NO, the gene content of *w*VitA is closer to that of *w*OneA1 than *w*VitB, as expected by their supergroup affiliations and indicating that there is no rampant recombination between the wVitA and wVitB at genome-wide level.

**Table 2.** Number of IS in *w*One, *w*VitA and *w*VitB.

| **Family** | **Number in *w*One** | **Number in *w*VitA** | **Number in *w*VitB** |
| --- | --- | --- | --- |
| IS3 | 0 | 0 | 0 |
| IS4 | 0 | 1 | 0 |
| IS5 | 1 | 1 | 1 |
| IS5/IS1182 | 0 | 1 | 0 |
| IS6 | 0 | 0 | 2 |
| IS66 | 0 | 2 | 0 |
| IS110 | 2 | 2 | 0 |
| IS200/IS605 | 2 | 0 | 1 |
| IS256 | 0 | 1 | 2 |
| IS481 | 0 | 0 | 2 |
| IS630 | 0 | 0 | 2 |
| IS982 | 0 | 1 | 0 |

### Phylogenomic analysis of 34 *Wolbachia* genomes revealed the evolutionary relationship and potential recombination events among strains

We compared *w*OneA1 to 33 other sequenced *Wolbachia* genomes (Table S1), which include 15 A-group and 12 B-group strains from diverse host species. Single-gene ortholog clusters were generated using the procedure described in the Methods, and 211 single-gene ortholog clusters (listed in Supplemental Data S1) were identified that are shared between *w*OneA1 and the 33 other *Wolbachia* genomes. This is a smaller set than the 496 *Wolbachia* gene orthologs detected in (Lindsey, et al. 2016) for 16 *Wolbachia* strains, but ours includes a larger strain set (34 *Wolbachia* strains) and we restricted our analysis to single copy orthologs across the genomes. Based on the coding sequences of this core gene set, a Maximum Likelihood (ML) phylogenetic tree of 34 *Wolbachia* genomes confirmed the separation of different supergroups A (*w*Suzi, *w*Spc, *w*Ri, *w*Ha, *w*Au, *w*Mel, *w*MelPop, *w*Gmm, *w*Uni, *w*DacA, *w*Nfe, *w*Npa, *w*Nfla, *w*Nleu, *w*VitA, *w*OneA), B (*w*Con, *w*AlbB, *w*Stri, *w*Di, *w*No, *w*Tpre, *w*DacB, *w*VitB, Ob_Wba, *w*Bol1, *w*Pip_Mol, *w*Pip), C (*w*Oo, *w*Ov), D (*w*Bm, *w*Wb), E (*w*Fol) and F (*w*Cle) with 100% bootstrap support (Figure 4, Supplemental Data S2 and S3). Noticeably, *w*OneA1 is more closely related to a subset of A-*Wolbachia* found in *Drosophila* (*w*Ha, *w*Ri, *w*Spc and *w*Suzi) than to *w*VitA and *w*Uni in parasitoid wasps. This pattern was previously observed using MLST by five genes in *Wolbachia* (Raychoudhury, et al. 2009), but is now supported by a much larger data set. Our genomic analyses also supported extensive horizontal movement of *Wolbachia* strains between divergent host species. A ML phylogenetic tree of protein sequences from these core genes was also constructed with RAxML. The protein ML phylogenetic tree is highly similar to the coding sequence ML tree with some minor differences (Figure S1, Supplemental Data S4 and S5).

**Figure 4.**
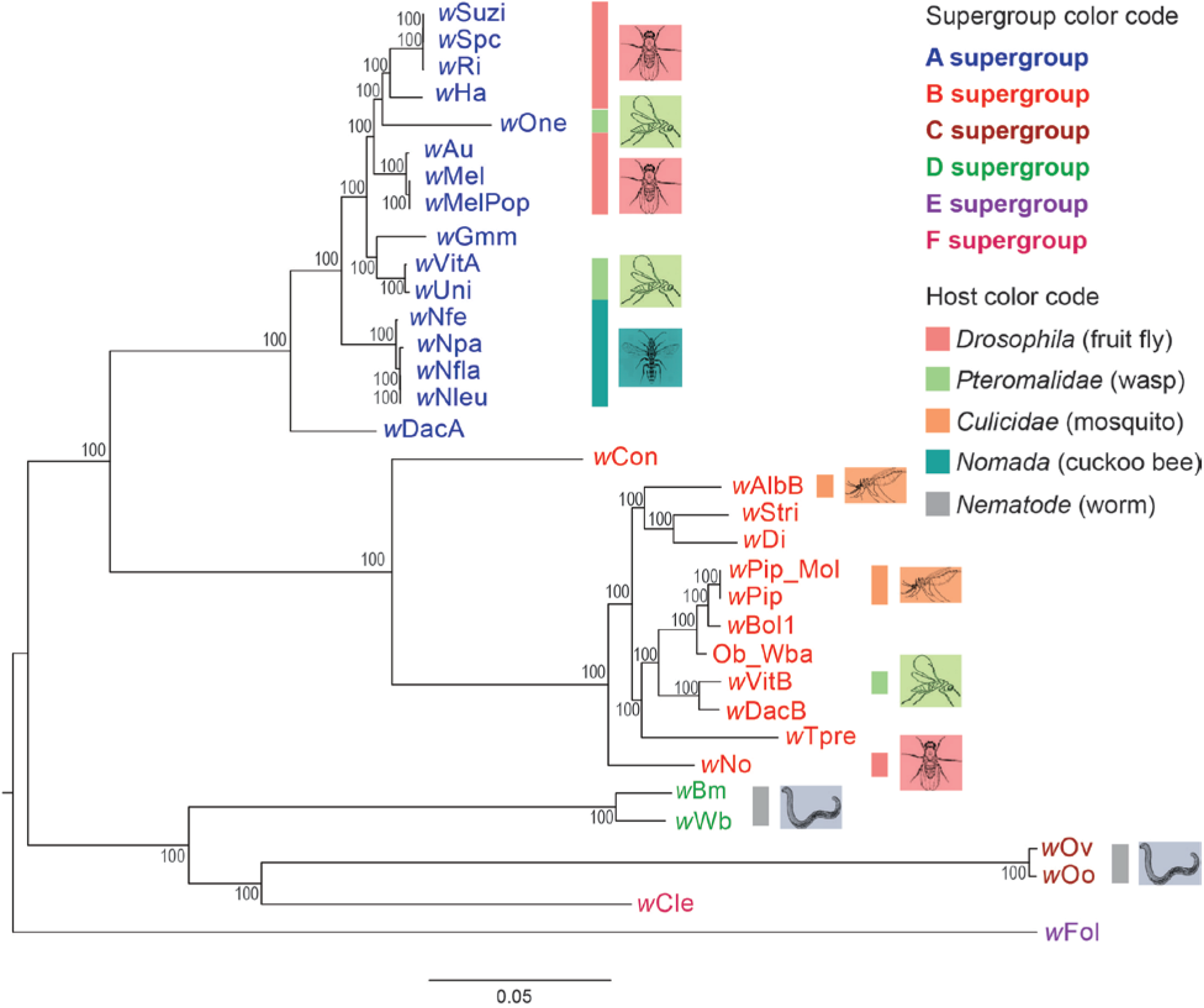
Phylogenomic relationships of 34 *Wolbachia* strains. The phylogenetic tree was constructed using Maximum Likelihood method from a concatenated coding sequence alignment of 211 single-copy orthologous genes with RAxML. Numbers on the branches represent the support from 1000 bootstrap replicates. The supergroup classifications (A-F) represent following the color code. The host taxonomic classifications for most of *Wolbachia* strains are shown in different color code.

We next examined genomes for potential lateral gene transfers among the core gene set. Single gene trees of coding sequences from the core gene set were constructed to check for supergroup level consistency. For 205 of the 211 trees, the separation among the A to F supergroups was consistent, with slight rearrangement for some *Wolbachia* strains within each supergroup. However, six trees are mixed among different supergroups, presumably due to lateral gene transfer or recombination events between strains (Figures 5–8, Supplemental Data S6-S11); the genes are hypothetical protein WONE_01840, cytochrome c oxidase subunit II (coxB), hypothetical protein WONE_04820, NADH-quinone oxidoreductase subunit C (nuoC), molecular chaperone (DnaK), and arginine-tRNA ligase (argS).

**Figure 5.**
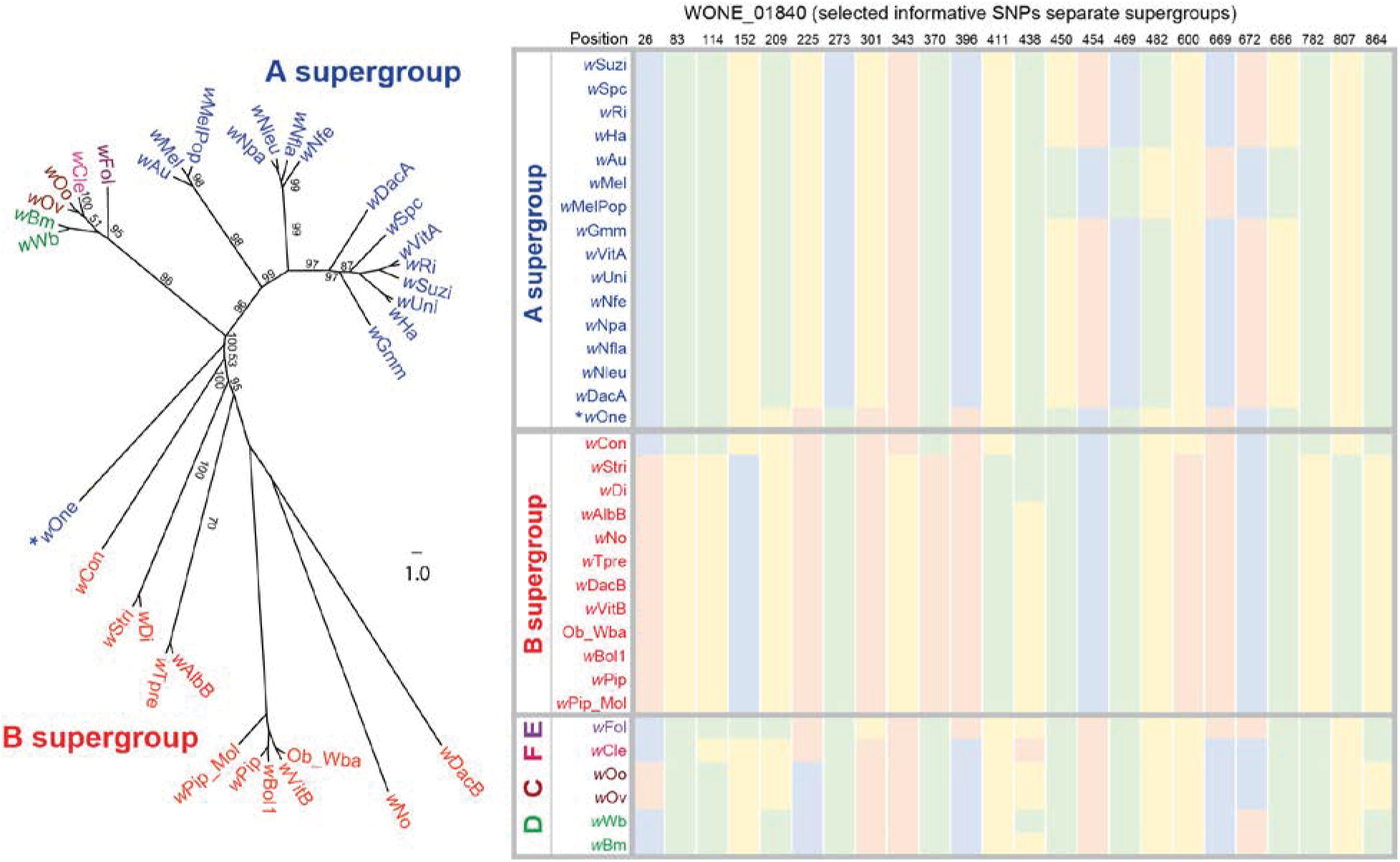
RAxML tree revealed lateral gene transfer between *w*One and B-*Wolbachia*. The supergroup classifications follow the color code in Figure 4. Nucleotides at selected positions are shown in the right panels (green:A; blue: C; yellow: G; pink: T). Hypothetical protein WONE_01840 from *w*One (A supergroup) clusters with B-*Wolbachia*.

**Figure 6.**
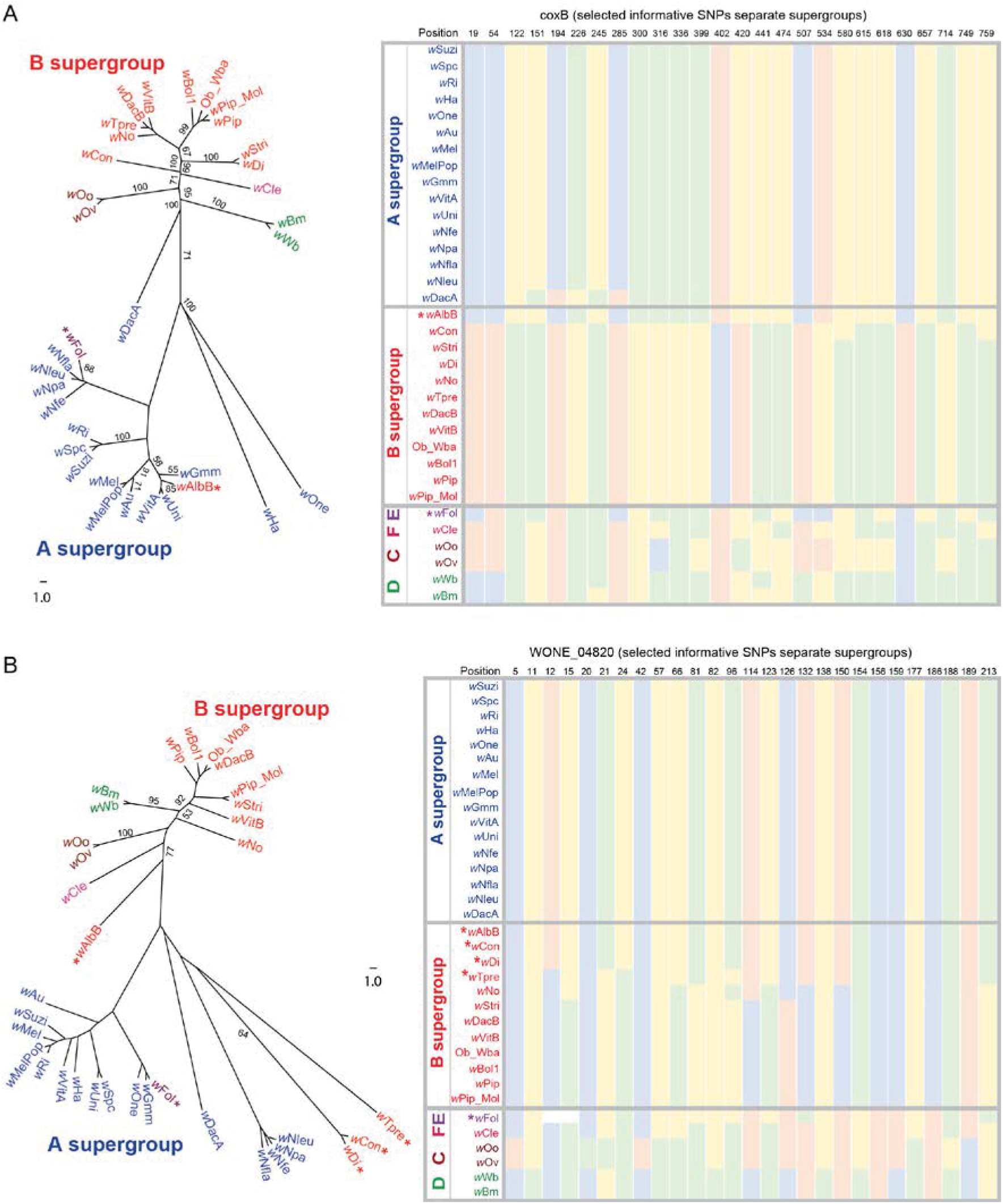
RAxML trees revealed lateral gene transfer between A-*Wolbachia* and B/E-*Wolbachia* in coxB and WONE_04820 genes. The supergroup classifications follow the color code in earlier figures. Nucleotides at selected positions are shown in the right panels. (A) In ML tree of coxB, *w*AlbB (B supergroup) and *w*Fol (E supergroup) cluster with A-*Wolbachia*, respectively; (B) In ML tree of hypothetical protein WONE_04820, *w*Con, *w*Di and *w*Tpre and *w*AlbB (B supergroup) cluster with A-*Wolbachia*, and *w*Fol (E supergroup) clusters with A-*Wolbachia*.

**Figure 7.**
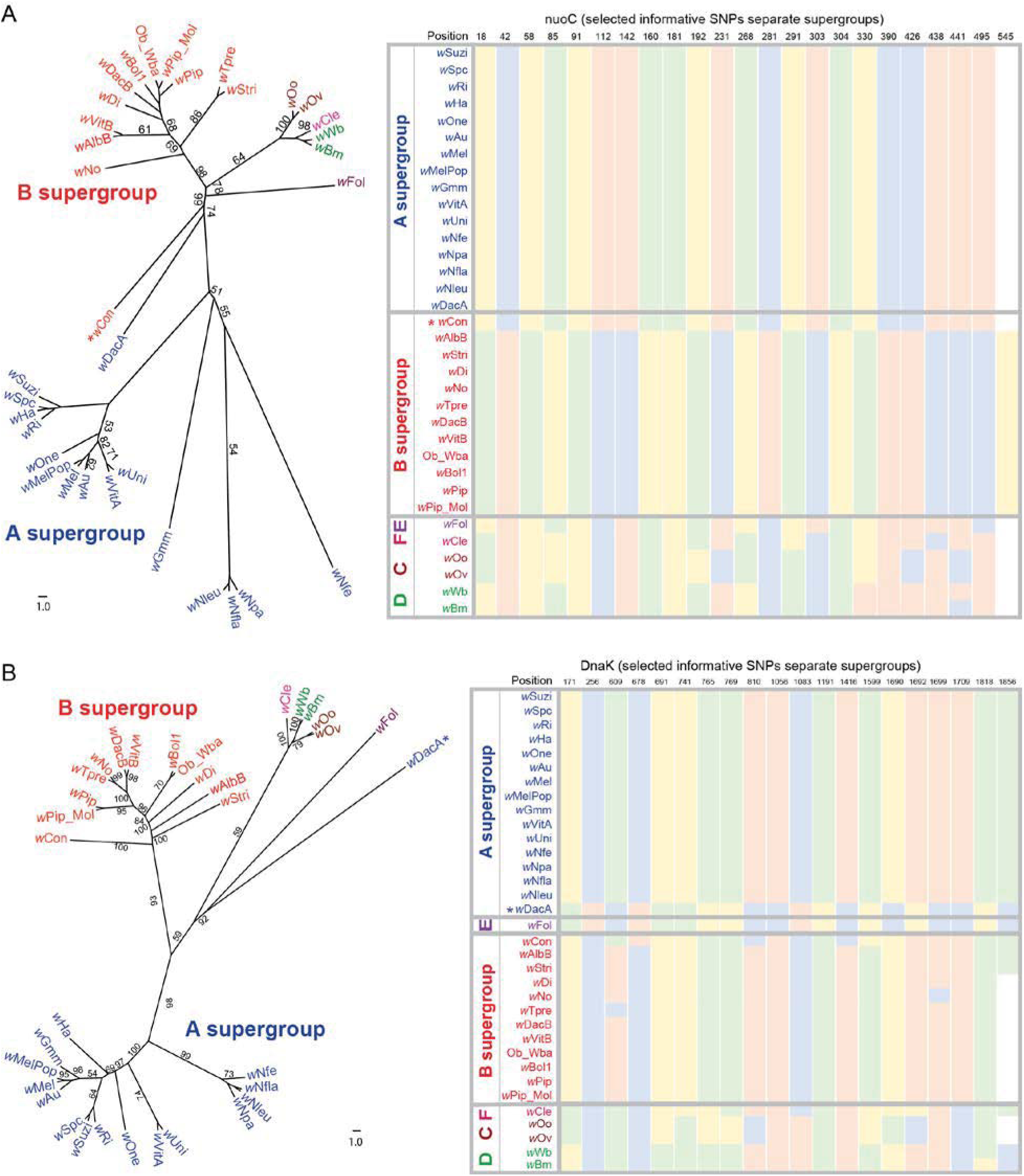
RAxML trees revealed lateral gene transfer between A-*Wolbachia* and B/E-*Wolbachia* in nuoC and DnaK genes. The supergroup classifications follow the color code in earlier figures. Nucleotides at selected positions are shown in the right panels. (A) in ML tree of nuoC, *w*Con (B supergroup) clusters with A-*Wolbachia*. (B) in ML tree of DnaK, *w*DacA (A supergroup) clusters with *w*Fol (E supergroup).

**Figure 8.**
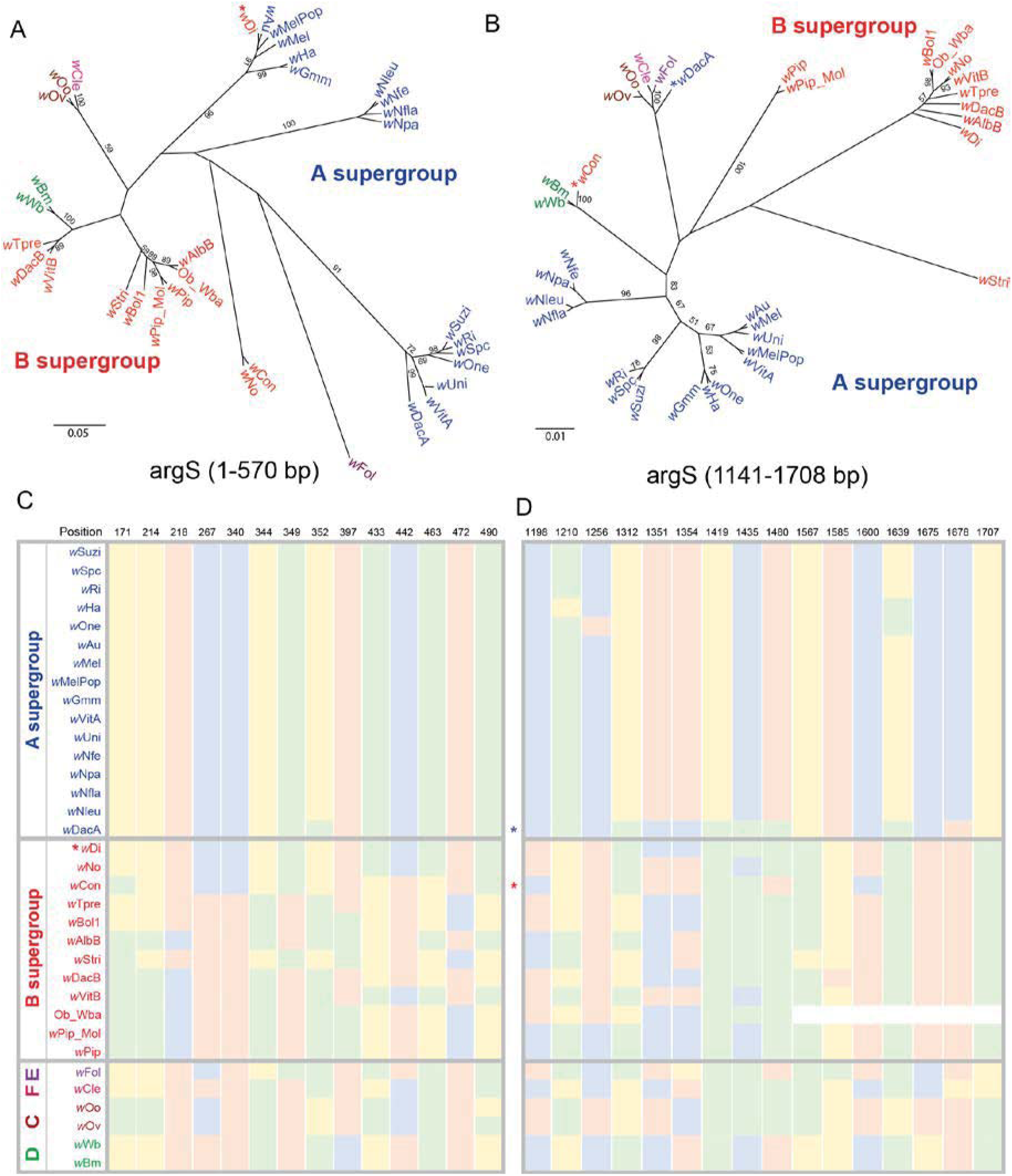
Distinct tree structure difference revealed in argS gene. The supergroup classifications follow the color code in earlier figures. (A) Tree structure of argS (from starting site to 570 bp) revealed that *w*Di (B supergroup) clusters with A-*Wolbachia*; (B) Tree structure of argS (from 1141 bp to stop site) revealed that *w*DacA (A supergroup) clusters with E-*Wolbachia* and *w*Con clusters with D-*Wolbachia*, indicating intragenic recombination events; (C) Nucleotides at selected positions (1-570 bp in argS) supported the tree topology in (A); (D) Nucleotides at selected positions (1141-1708 bp in argS) supported the tree topology in (B).

Among these six single gene trees, one tree showed evidence of a recombination event between B-*Wolbachia* and A/D-*Wolbachia* in gene argS (Figure 8, Supplemental Data S11). Four trees revealed some mix groupings between A-*Wolbachia* and B-*Wolbachia* in hypothetical protein WONE_01840 (Figure 5, Supplemental Data S6), coxB (Figure 6A, Supplemental Data S7), hypothetical protein WONE_04820 (Figure 6B, Supplemental Data S8), and nuoC (Figure 7A, Supplemental Data S9), which suggest a lateral transfer. Also, three trees grouped A-*Wolbachia* and the E-*Wolbachia w*Fol together, including *w*Fol clustered with A-*Wolbachia* in coxB (Figure 6A) and WONE_04820 (Figure 6B), and A-*Wolbachia w*DacA clustered with *w*Fol in DnaK (Figure 7B, Supplemental Data S10) and argS (Figure 8B). The recombination between A and B supergroups in gene coxB was reported by a previous study of 6 *Wolbachia* strains (Ellegaard, et al. 2013), and the remaining identified between-supergroup recombination events are novel findings in our study. Taken together, 97% of the single copy orthologs agree with the supergroup classification in *Wolbachia*, with a few cases of likely recombination or lateral transfer events between *Wolbachia* strains of different supergroups. The finding also indicates that these recombination events involve relatively small regions, rather than large recombination events involving many genes. The frequent gene order rearrangements observed in *Wolbachia* may make larger recombination tracks between supergroups less successful, as they are more likely to involve vital gene losses due to lack of synteny.

### Phylogenetic analysis of MLST genes of *Wolbachia* in *Nasonia*

Previous research indicated that NO contains three *Wolbachia* strains, A1, A2 and B, which were acquired through a hybrid introgression from NG (Raychoudhury, et al. 2009). However, the genome sequence for the current *N. oneida* strain indicates presence of only one A-group strain, even though the same insect strain is present in both studies. This difference is likely due to stochastic loss of two strains during laboratory culturing. This is known to happen in *Nasonia*, particularly when the strains are passed through winter larval diapause (Perrot-Minnot, et al. 1996), which involves storage under refrigeration for up to 1.5 years after larval diapause induction. To determine which strain (A1 or A2) was identified and assembled in our study, phylogenetic analysis was conducted with all *Wolbachia* strains in *Nasonia* using the MLST gene approach (Baldo, et al. 2006b; Paraskevopoulos, et al. 2006). The phylogenetic trees were constructed for all five MLST genes (Figure 9). *w*One MLST genes grouped with A1 strains of other species. The closest branch is the corresponding *w*NgirA1 ortholog for all five MLST genes, indicating that the assembled *w*One is the strain A1. All phylogenetic trees of five MLST genes supported the conclusion that the identified *w*One in our study belongs to the A1 strain of NO. The MLST gene sequence of *Wolbachia* in NO are the same as the corresponding one of *Wolbachia* in NG (Raychoudhury, et al. 2009). Our results are consistent with the previous findings for all five MLST genes in *w*OneA1.

**Figure 9.**
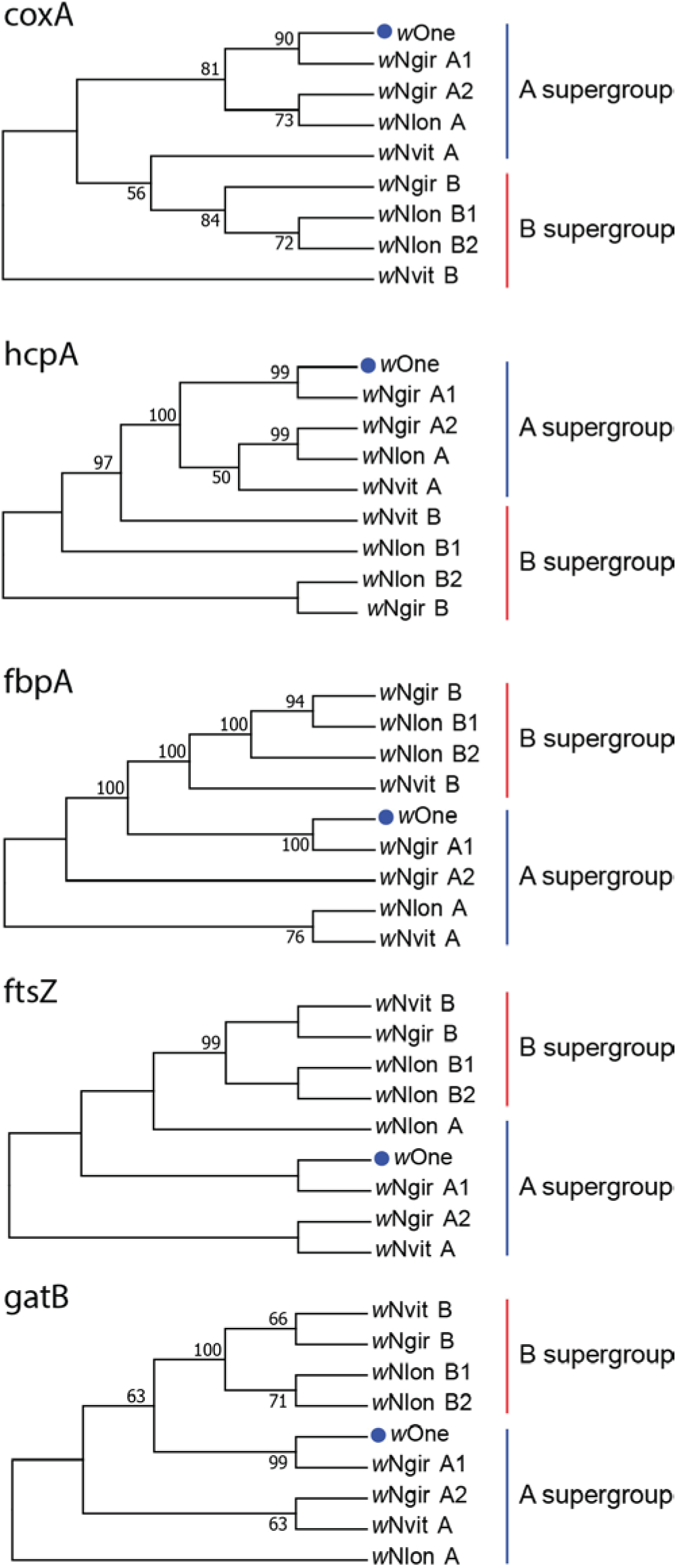
Phylogenetic tree of *Wolbachia* in *Nasonia* with Maximum Likelihood method based on five MLST genes. The bootstrap consensus tree inferred from 1000 replicates. A and B supergroups were clustered into two groups in trees of all five MLST genes.

### Loss of A2 and B *Wolbachia* in the assembled *N*. *oneida* strain

To determine whether there are other *Wolbachia* strains in NO, we aligned the *w*One sequencing reads to NG MLST genes. No reads were mapped to *w*NgirB MLST genes, suggesting the B supergroup is absent in our NO strain. For *w*NgirA1 MLST genes, the average coverage is 30X which are close to the coverage of *w*One scaffolds. *w*NgirA2 MLST genes only have multiple mapped reads to both *w*NgirA1 and *w*NgirA2, which is not informative to determine the existence of the A2 strain. The informative SNPs in each of MLST genes between *w*NgirA1 and *w*NgirA2 were further checked for read counts to ensure the inability to detect of *w*OneA2. No read count was identified for A2 allele of all MLST genes, while all A1 alleles were supported by at least 30 read counts (Table S2).

Furthermore, strain typing of *Wolbachia* was performed on NO of our study and NO genomic DNA samples that are known to be infected with all three strains (A1, A2 and B), using independent allele-specific pyrosequencing approach. An A/G SNP in the coxA gene was used to separate B-*Wolbachia* from A-*Wolbachia* (A allele in A1/A2-*Wolbachia* and G allele in B-*Wolbachia*, Figure S2A). In gatB gene, a C/T SNP can distinguish A1-*Wolbachia* allele from A2/B-*Wolbachia* (Figure S2B). The pyrosequencing results confirmed the lack of A2 and B strains in the genome assembled NONY strain. All three *Wolbachia* infections (A1, A2 and B) were successfully identified in the CAR262L strain DNA samples (Figure S2 and Table S3).

### Single Nucleotide Polymorphisms (SNPs) likely represent gene duplications within the *w*OneA1 genome are enriched in transmembrane transporter genes

A total of 68 high-quality SNPs was called using *w*OneA1 genome alignments (Figure 10). The alternative allele frequency of these identified SNPs ranges from 0.21 to 0.62 (Figure S3). All identified SNPs were shown in the circular view of *w*One genome (Figure 10). Read alignments for SNP positions were visualized by IGV (two examples were listed in Figure 10). For most cases, the SNPs are clustered in regions with higher (2-3 fold) coverage than the rest of the *w*OneA1 genome, suggesting gene duplication might be the cause of polymorphisms in *w*One genome (Figure 10). Among these identified SNPs, 27 were found to be located within gene regions (Table S4). They likely did not assemble as unique duplications in the genome due to sequence similarity. For instance, we identified multiple SNPs located in the region of WONE_08300 gene in SCAFFOLD10, which is phage-related baseplate assembly protein J (Figure 10A). Reads alignment in IGV indicated that most SNPs were linked, and we can manually assemble these reads into two homologous genes (Figure 10B and 10C). The protein sequence of the alterative assembly has 92% identity with baseplate assembly protein J. However, long read technology would be needed to resolve their status as duplications. BLAST2GO (Conesa, et al. 2005) analysis identified the transmembrane transporter activity molecular function GO term was significantly enriched (Figure S4) among genes containing SNPs.

**Figure 10.**
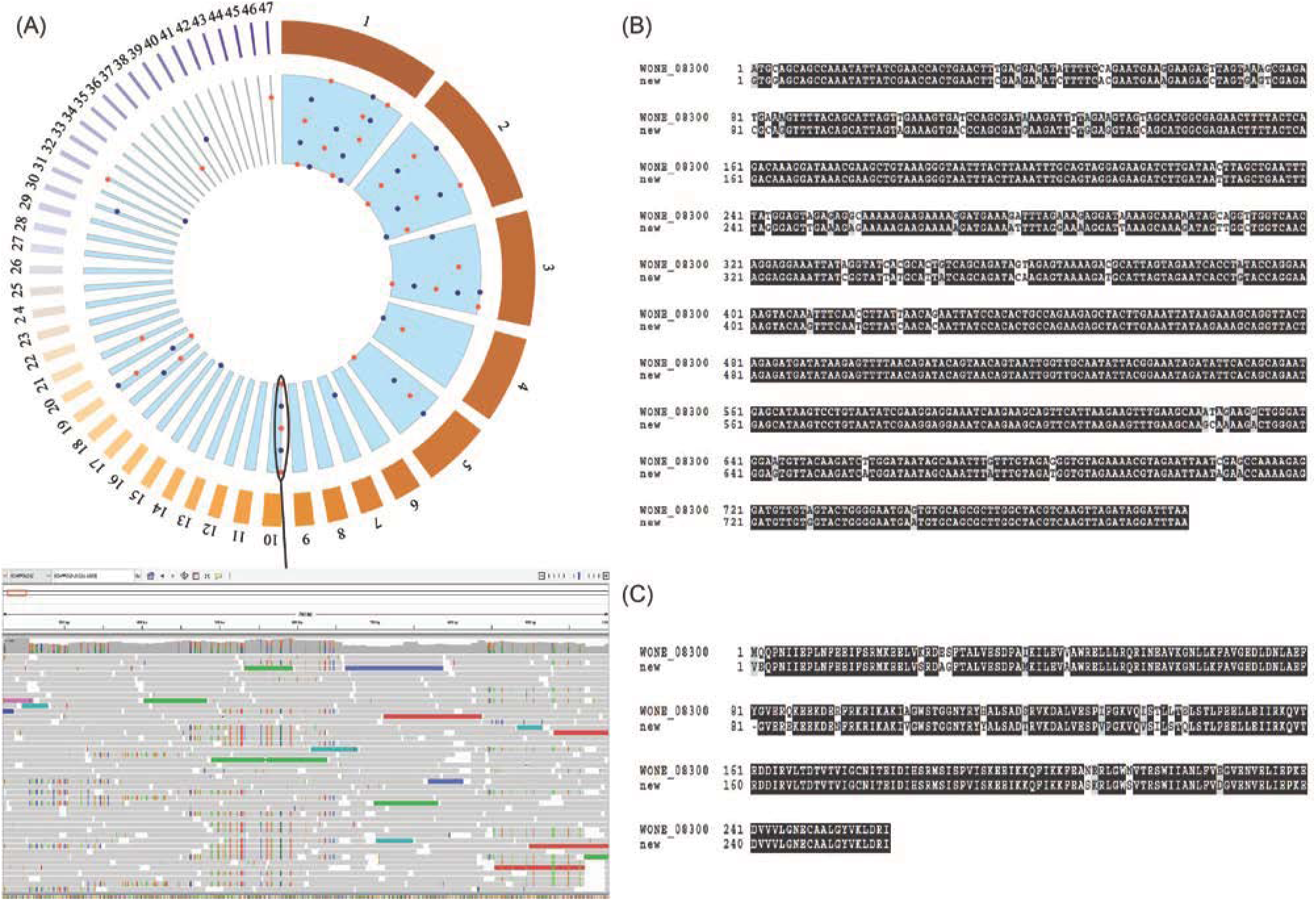
Polymorphic SNPs in *w*One genome. (A) Circular view of *w*One genome with the blue and orange spots indicate the identified SNP positions. Read alignments for SNPs located in WONE_08300 gene region in Scaffold10 was shown in IGV screenshots, suggesting these polymorphisms result from gene duplications in *w*One genome. (B) Nucleotide sequence alignment between new assembly from reads with SNPs and WONE_08300 gene. (C) Protein sequence alignment between new assembly from reads with SNPs and WONE_08300 gene.

### Concordance of MLST genes and whole genome divergence

The MLST system has been variously used for strain typing of *Wolbachia*, identification of related strains, recombination within genes (e.g. the *wsp* locus) and for phylogenetic inferences among strains. Recently, reliability of the MLST system has been criticize (Bleidorn and Gerth 2018) as unreliable, with whole genome sequencing to be preferred. Although whole genome data sets would always be desirable, we undertook to compare genetic divergence based the MLST to our set of 211 genes in 34 different *Wolbachia* strains. The MLST performed very well in both identifying closely related strains and in genetic divergence among strains compared to the genome wide data set. The correlation coefficient (rho) of estimated evolutionary divergence using core gene set and gatB, fbpA, hcpA, coxA, ftsZ is 0.96, 0.9, 0.97, 0.92 and 0.97, respectively with *P-value* < 2.2 × 10^−16^ (Table 3, Figure S5, Supplemental Data S12). Eventually, whole genome data sets will supplant the MLST system. However, with over 1900 isolates in the *Wolbachia* MLST database, this will likely take some time, and until then, MLST remains a reliable method for identifying closely related *Wolbachia* strains and their host associations.

**Table 3.** Estimates of evolutionary divergence between *Wolbachia* species using core gene set and five MLST genes.

| **Correlation coefficient (rho)** | **core gene set** | **gatB** | **fbpA** | **hcpA** | **coxA** | **ftsZ\*** |
| --- | --- | --- | --- | --- | --- | --- |
| **core gene set** | 1 | 0.96 | 0.90 | 0.97 | 0.92 | 0.97 |
| **gatB** |  | 1 | 0.86 | 0.91 | 0.90 | 0.94 |
| **fbpA** |  |  | 1 | 0.87 | 0.84 | 0.92 |
| **hcpA** |  |  |  | 1 | 0.89 | 0.96 |
| **coxA** |  |  |  |  | 1 | 0.92 |
| **ftsZ\*** |  |  |  |  |  | 1 |

\*Estimates of evolutionary divergence using ftsZ gene were only conducted among 31 *Wolbachia* species excluding wBm, wWb and wCon, because of the inability to correctly annotate ftsZ in these 3 species.

## Discussion

### Assembly of a prokaryotic genome in an insect species using linked reads technology

Here we report the first *Wolbachia* genome assembled from host sequences using 10X Genomics linked-reads technology. Although many *Wolbachia* genomes have been assembled since the first *w*Mel genome paper was published, the difficulty in purifying *Wolbachia* DNA from the host is still a limiting factor for the genomic studies of *Wolbachia*. Due to its intracellular lifestyle and inability of media culture, the purification of *Wolbachia* DNA from the host sample is time-consuming and sometimes impossible to obtain sufficient quantity without contamination from the host nuclear and mitochondrial genomes (Table S1). Three major methods have been applied to purify the *Wolbachia* genomic DNA from the host DNA: 1) selection of *Wolbachia* enriched materials for DNA extraction, such as host ovaries (Brelsfoard, et al. 2014) or *Wolbachia* infected cell lines (Mavingui, et al. 2012); 2) use of different filter purification methods including pulsed-field gels (Wu, et al. 2004), various gradient gels (Duplouy, et al. 2013; Klasson, et al. 2009), or filter columns (Newton, et al. 2016); 3) Multiple-displacement amplification (MDA) for *Wolbachia* DNA enrichment and amplification (Ellegaard, et al. 2013; Mavingui, et al. 2012). Despite the efforts on the purification of *Wolbachia* genomic DNA, only 90% purification could be achieved. Alternative methods have been applied to solve the contamination problem. For example, in *w*Bm genome project, *Wolbachia* BACs were selected from the host BAC library (Foster, et al. 2005). For the *w*VitB genome project, a high-density tiled oligonucleotide array was developed to enrich for *Wolbachia* gDNA (Kent, et al. 2011).

If no prior knowledge is available about the presence of specific microbes, sequencing without purification is preferred to identify other intracellular symbionts, as well as characterizing bacterial species in insect gut microbiota at the whole-genome level. Other studies have extracted *Wolbachia* reads from the host whole genome sequence dataset, and then align to the reference genome of the closely related *Wolbachia* strains, or perform *de novo* assembly using the filtered reads (Chung, et al. 2017; Darby, et al. 2012; Lindsey, et al. 2016; Saha, et al. 2012; Siozios, et al. 2013). Here we perform *de novo* assembly of the host genome and *Wolbachia* genome using the 10X Genomics linked-reads technology. The *Wolbachia* DNA fragments were labeled with unique 10X barcodes, therefore they are much less likely to be misassembled into the host genome scaffolds. In this study, the microbe and the host have a 10-fold coverage difference, allowing accurate identification of the bacterial scaffolds. Therefore, the *Wolbachia* genome assembled with 10X linked reads was of good quality with no contamination of host nuclear and mitochondrial DNA. As the cost of PacBio sequencing decreases, the long-read platforms would be better for symbionts genome assembly, unless the bacterial reads are much lower than the host DNA.

### Lateral gene transfer and recombination events among *Wolbachia* genomes

The phylogenomic analysis of 34 *Wolbachia* genomes in our study is the most comprehensive phylogenomic and evolutionary analysis conducted in *Wolbachia* strains to date. By including almost all available *Wolbachia* genomes in NCBI, we confirmed at the genome level that these *Wolbachia* strains group into distinct clusters (A, B, C, D, E, F supergroups) and different *Wolbachia* co-infected in the same host kept the strain boundary (Ellegaard, et al. 2013). 205 of the 211 single gene trees are consistent with the strain tree. Six genes trees have major rearrangements among *Wolbachia* groups (Figures 5–8), indicating potential recombination or lateral gene transfer events between strains. We estimated that the homologous recombination occurred in at least 3% of the core genes in the *Wolbachia* genomes, and recombination may be one of the evolutionary forces shaping the *Wolbachia* genomes.

The six genes with distinct tree structure differences from the consensus *Wolbachia* tree include coxB, nuoC, DnaK, argS, hypothetical proteins WONE_01840 and WONE_04820. Gene coxB and nuoC are both involved in the electron transport chain, which might also interact with hypothetical proteins WONE_01840 and WONE_04820. The functions of hypothetical proteins WONE_01840 and WONE_04820 in *Wolbachia* are still unclear. However, the potential recombination or lateral gene transfer events identified in these genes might suggest that their maintenance could be under positive selection.

LGT events among A and B *Wolbachia* supergroups have been documented in previous studies, and we identified a few addition cases through the phylogenomic analysis among 34 sequenced genomes. Interesting we also discovered LGT events between A and E supergroups, which was not known previously. The E group *Wolbachia* was found in Collembola, or the springtails (Czarnetzki and Tebbe 2004; Fountain and Hopkin 2005; Vandekerckhove, et al. 1999). A recent study characterized the *Wolbachia* in 11 collembolan species, and found that nearly all are E group *Wolbachia* that are monophyletic, based on phylogenetic reconstruction using MLST genes (Ma, et al. 2017). Our genome analysis of the single collembolan *Wolbachia* genome reveals a number of candidate lateral gene transfer events, including intragenic recombination in argS between A, B, D and E, lateral gene transfer in coxB and DnaK, and in WONE_04820. Targeted sequencing of these genes in the additional collembolan species or additional genome sequences will help reveal the origins and directions of these lateral gene transfers. We further speculate that selective maintenance of such transfers could suggest a possible role in the *Wolbachia* function, such as parthenogenesis induction (Ma, et al. 2017).

It has been recently argued that MLST genotyping has little utility in phylogenetic analyses, and should be supplanted by genomic studies (Bleidorn and Gerth 2018). When the MLST system was developed, it was pointed out by the authors that the system would be most useful for identifying relatively closely related *Wolbachia*, due to potential recombination among more divergent strains (Baldo and Werren 2007). However, our comparison on genome sequence indicates that MLST typing is largely valid, both for supergroup identification and detection of closely related strains. Related *Wolbachia* based on MLST results are also genome-wide closely related. This suggests that, until *Wolbachia* genome sequencing becomes much less expensive and can be readily performed on single arthropods, that MLST will remain a useful tool for identification of strains, their relationships, and host affinities. Nevertheless, caution should be exercised due to some documented recombination events within MLST genes and among them (Raychoudhury, et al. 2009). Therefore, topologies should be compared among genes for evidence of discordance, rather than simply relying of phylogenetic reconstructions of concatenated sequences.

### Loss of A2 and B *Wolbachia* in *N. oneida* lab strains

The whole genome alignments between *w*One and *w*VitA (Figure 3) and the phylogenetic analysis of MLST genes (Figure 9) indicated the identified strain in our study is belong to A1 supergroup. The results indicate either A2 and B are present in an extremely low level so that they cannot be detected at the current sequencing coverage, or their density is much less in our studied male NO adult samples. Subsequent allele-specific pyrosequencing validation experiments confirmed the absence of A2 and B-type *Wolbachia* infections. In NO DNA samples from a recently collected field strain, we estimate that A1 is the dominate strain and accounts for 55% of the total infection, 40% of the infection came from the B strain and only 5% from A2 strain (Figure S2). The absence of A2 and B *Wolbachia* in the lab NO strain is likely due to stochastic loss during laboratory maintenance and diapause.

### Evolution of the *Wolbachia* genome in the *Nasonia* genus

The draft genome of *w*OneA1 is comprised of 47 scaffolds with a total length of 1.29 Gb. The total size of *w*One draft genome is relatively longer when comparing to the other two *Nasonia*-associated *Wolbachia w*VitA and *w*VitB. Most of the genomic regions in *w*OneA1 draft genome aligned well with their corresponding regions in *w*VitA with several rearrangements, indicating syntenic conservation between these two strains (Figure 3A). However, there are more structural differences between A-*Wolbachia* and B-*Wolbachia*, which were supported by the whole genome alignments between A-*Wolbachia* (*w*One and *w*VitA) and B-*Wolbachia w*VitB (Figure 3B and C). The same pattern was observed when comparing gene contents among these three *Wolbachia* in *Nasonia*, *w*OneA1 and *w*VitA shared more orthologous genes (Figure 3D). The results indicate that A and B *Wolbachia* retain their genetic differences even when they infect the same host, which suggests that recombination among them is not common, with the exception of phage related genes (Bordenstein and Wernegreen 2004).

We have sufficient sequence coverage in *w*One genome to identify segregating SNPs, but all the candidate SNPs are located in regions with elevated sequencing depth (Figure 10), suggesting they are fixed differences in recently duplicated genes with multiple copies rather than segregating SNPs. This is consistent with the severe bottleneck due to the maternal transmission of *Wolbachia* through the egg, which is extremely hard to maintain segregating SNPs through balancing selection. These potential newly duplicated genes are enriched for transmembrane transporter function. Due to the intercellular lifestyle, the membrane proteins are critical for *Wolbachia* infection, nutrient uptake and other interactions with the host cells. These duplication events may provide advantage over the A2 and the B strains.

## Methods

### Sample collection, genomic DNA extraction, 10X Genomic library preparation and genome sequencing

Genomic DNA sample was extracted from 24-hour male adults of the *N. oneida* NONY strain. MagAttract HMW DNA Mini Kit (Qiagen, MD) was used to isolate high molecular weight genomic DNA. The quality of extracted gDNA was checked on a Qubit 3.0 Fluorometer (Thermo Fisher Scientific, USA). The size distribution of the extracted gDNA was accessed using the genomic DNA kit on Agilent TapeStation 4200 (Agilent technologies, CA). A 10X Genomic library was constructed by using the Chromium Genome Reagent Kits v2 on 10X Chromium Controller (10X Genomics Inc., CA). Chromium i7 Sample Index was used as library barcode. Post library construction quality control was accessed with Qubit 3.0 Fluorometer and Agilent TapeStation 4200. The constructed 10X genomic library was sequenced on a HiSeq X sequencer at the Genomic Services Lab at the HudsonAlpha Institute for Biotechnology.

### Genome assembly and annotation of *w*One genome using linked reads

The *N. oneida* genome was assembled using the Supernova 2.1.1 assembler (Weisenfeld, et al. 2017) with all 10X linked reads. The following steps were conducted to identify *w*One scaffolds in the *N. oneida* assembly (Figure 1): 1) all sequencing reads were aligned to the *N. oneida* assembly to calculate the average and median coverage for each scaffold in the assembly; 2) *N. oneida* scaffolds are aligned to the bacterial sequence database using BLAT version 3.5 (Kent 2002) to determine the percent of sequence identity to known *Wolbachia* sequences; 3) assign the scaffolds to *w*One genome. A scaffold was assigned to *w*One genome if it has at least 20% sequence identity with known *Wolbachia* sequences and a median coverage around 60X and GC content around 0.35. The genome completeness was further evaluated by checkM (Parks, et al. 2015) with default settings and BUSCO (Seppey, et al. 2019) comparing to bacteria database. Gene annotation was conducted using DFAST prokaryotic genome annotation pipeline (Tanizawa, et al. 2018) with few manual correction based on other *Wolbachia* gene models. tRNA genes were predicted by tRNAscan_SE (Lowe and Eddy 1997).

### Comparative analysis of *Wolbachia* genomes in the *Nasonia* genus

To compare the genome structure among three sequenced *Wolbachia* genomes in *Nasonia*, we first conducted whole genome alignment of *w*One, *w*VitA (GenBank accession GCA_001983615.1) and *w*VitB (GenBank accession GCA_000204545.1) (Kent, et al. 2011) genomes using NUCmer in the MUMmer program suite with default parameter settings (Kurtz, et al. 2004). The pairwise alignments (match length longer than 500bp) were visualized using Mummerplot (Kurtz, et al. 2004). Orthologous gene sets between *w*One and two other *Wolbachia* in *Nasonia* were generated based on reciprocal best hits using BLAST with an E-value cutoff 10^−5^. 32 genes in *w*One genome were excluded in this analysis as the gene orthologies are unclear when comparing to *w*VitA and *w*VitB.

### Phylogenomic analysis of 34 genome sequenced *Wolbachia* strains

To examine the phylogeny of *Wolbachia* at the genome level, we conducted phylogenomic analysis using *w*One and 33 other sequenced *Wolbachia* genomes (GenBank accession numbers listed in Table S1), including *w*Mel (Wu, et al. 2004), *w*Bm (Foster, et al. 2005), *w*Oo (Darby, et al. 2012), *w*Pip (Klasson, et al. 2008), *w*Ri (Klasson, et al. 2009), *w*VitB (Kent, et al. 2011), *w*Bol1 (Duplouy, et al. 2013), *w*Ha (Ellegaard, et al. 2013), *w*No (Ellegaard, et al. 2013), *w*Gmm (Brelsfoard, et al. 2014), *w*AlbB (Mavingui, et al. 2012), *w*VitA (Newton, et al. 2016), *w*Di (Saha, et al. 2012), *w*Suzi (Siozios, et al. 2013), *w*Tpre (Lindsey, et al. 2016), *w*Wb (Chung, et al. 2017; Desjardins, et al. 2013), *w*Ov (Desjardins, et al. 2013), *w*Pip_Mol (Pinto, et al. 2013), *w*Au (Sutton, et al. 2014), Ob_Wba (Derks, et al. 2015), *w*Cle (Nikoh, et al. 2014), *w*Fol (Faddeeva-Vakhrusheva, et al. 2017), *w*Con (Badawi, et al. 2018), *w*DacA (Ramirez-Puebla, et al. 2016), wDacB (Ramirez-Puebla, et al. 2016), *w*MelPop (Woolfit, et al. 2013), *w*Nfe (Gerth and Bleidorn 2017), *w*Npa (Gerth and Bleidorn 2017), *w*Nleu (Gerth and Bleidorn 2017), *w*Nfla (Gerth and Bleidorn 2017), *w*Spc (Conner, et al. 2017) and *w*Stri.

Homologous genes and ortholog clusters among all 34 *Wolbachia* genomes were determined by using OrthoFinder (Emms and Kelly 2015) with default settings. 211 core single-copy genes were identified for the subsequent analysis, their accession numbers are listed in Supplemental Table S5. The 211 core single-copy genes shared in all 34 *Wolbachia* genomes were aligned with MAFFT (Katoh and Standley 2014) at both nucleotide and protein sequence level. These single-gene alignments were concatenated into one alignment to use in the subsequent phylogenetic analysis. A Maximum Likelihood (ML) tree was constructed with the GTRGAMMA model and 1000 bootstrap replicates by RAxML v8.2 (Stamatakis 2014) using the concatenated coding sequence alignment of the core gene set. Similarly, the single gene trees for 211 core genes were generated by RAxML v8.2 (Stamatakis 2014) to check the consistency of supergroup classification. In addition, for phylogenetic analysis of protein sequences from the core gene set, the best-fit model of protein evolution was searched by ProtTest 3 (Darriba, et al. 2011). The final ML phylogenetic tree was inferred by using RAxML v8.2 (Stamatakis 2014) with the FLU protein model (best fit model identified by ProtTest 3) and 1000 rapid bootstrap replicates. All sequence alignment and tree files have been submitted and made publicly available through Dryad (doi:10.5061/dryad.kg87554).

### Phylogenetic analysis of *Wolbachia* in *Nasonia* using MLST genes

The five MLST (Multi Locus Sequence Typing) genes (Baldo, et al. 2006b; Jolley and Maiden 2010) were examined to further characterize the phylogenetic relationships of *Wolbachia* strains in *Nasonia*. These genes include *gatB* (aspartyl/glutamyl-tRNA (Gln) amidotransferase, subunit B), *coxA* (cytochrome c oxidase, subunit I), *hcpA* (conserved hypothetical protein), *ftsZ* (cell division protein) and *fbpA* (fructose-bisphosphate aldolase). The *w*One MLST genes were identified on five different genome scaffolds, including *coxA* on SCAFFOLD17, *gatB* on SCAFFOLD28, *hcpA* on SCAFFOLD47, *ftsZ* on SCAFFOLD73 and *fbpA* in SCAFFOLD76. MLST gene sequences of the following strains in different hosts were included in the analysis: *w*NvitA, *w*NvitB in N. *vitripennis*; *w*NgirA1, *w*NgirA2, *w*NgirB in N. *giraulti*; *w*NlonA, *w*NlonBl, *w*NlonB2 in N. *longicornis* (Raychoudhury, et al. 2009).

Sequences of MLST genes from *Nasonia Wolbachia* strains were downloaded from the MLST database (Baldo, et al. 2006b). Multiple sequence alignments were generated using MUSCLE (MUltiple Sequence Comparison by Log-Expectation) with default parameters (Edgar 2004). Phylogenetic analysis was performed using the ML method in MEGA 7.0 software (Kumar, et al. 2016). Bootstrap tests with 1,000 replicates were used to evaluate the phylogenetic trees.

In addition, the pairwise evolutionary divergence distances between *Wolbachia* species were estimated with both the core gene set identified in this study and five MLST genes in 34 *Wolbachia* species by using the Maximum Composite Likelihood model (Tamura, et al. 2004) in MEGA7 (Kumar, et al. 2016). Estimates of evolutionary divergence using ftsZ gene were only conducted among 31 Wolbachia species excluding wBm, wWb and wCon, because of the inability to correctly annotate ftsZ in these 3 species. The correlation coefficient (rho) of estimated evolutionary divergences with the core gene set and MLST genes was calculated with Hmisc package (Harrell Jr and Harrell Jr 2019) in R.

Previous study indicated that sequences of all MLST genes in *w*One are the same as that in *w*Ngir (Raychoudhury, et al. 2009). Therefore, we used sequences of MLST genes in *w*NgirA2 and *w*NgirB to check if the *w*OneA2 and *w*OneB can be detected in the studied NO. We aligned the sequencing reads to the NG MLST reference sequences from three *Wolbachia* strains in with BWA (Li and Durbin 2009), calculated the coverage, and further examined the alignment of each gene in Integrative Genomics Viewer (IGV) (Thorvaldsdottir, et al. 2013).

### Strain-typing of *Wolbachia* in *Nasonia* with MLST genes using pyrosequencing

*Wolbachia* infection types were further checked in the genome sequenced NO in this study, and genomic DNA samples from a recently (Summer 2018) collected wild-type CAR262L strain using allele-specific pyrosequencing. Pyro PCR and sequencing primers were designed to target SNP positions in coxA and gatB genes in A1, A2 and B *Wolbachia* using PyroMark Assay Design 2.0 (Qiagen, USA). A complete list of primers sequences could be found in Table S3. The A/G SNP targeted in coxA can separate B-*Wolbachia* from A1/A2-*Wolbachia*, and the C/T SNP in gatB allowed us to distinguish A1-*Wolbachia* from A2/B-*Wolbachia*. Pyrosequencing was performed on a Pyromark Q48 Autoprep instrument (Qiagen, USA) using the PyroMark Q48 Advanced CpG Reagents (Qiagen, USA). Briefly, the target regions in coxA and gatB genes were PCR-amplified using the biotin-labeled forward primers and the reverse primers using template genomic DNA samples. Then, pyrosequencing was performed on a PyroMark Q48 Autoprep instrument (Qiagen, USA) using the corresponding sequencing primers by following manufacturer’s protocol. The results were analyzed with Pyromark Q48 Autoprep software (Qiagen, USA). Three technical replicates were performed for each sample.

### *de novo* SNP calling in *w*One genome

To identify segregating polymorphisms in the *w*One genome, all sequencing reads were aligned to *w*OneA1 genome using BWA software (Li and Durbin 2009). Initial SNP calling were performed using SAMtools (Li, et al. 2009). SNPs were further checked manually in IGV (Thorvaldsdottir, et al. 2013) to filtered out low-quality and problematic SNPs. A total of 68 high-quality SNPs was kept for the subsequent analysis. The identified SNPs were shown in the circular view of *w*One genome using BioCircos (Cui, et al. 2016). Coding gene regions were extracted using BEDTools (Quinlan 2014) to annotate genic SNPs. SnpEff (Cingolani, et al. 2012) was used to predict the effects of these genetic variants. Gene Ontology (GO) annotation analysis was done on SNP containing genes using BLAST2GO with an E-value cutoff of 10^−5^ (Conesa, et al. 2005).

## Acknowledgements

This project is supported by an Auburn University Intramural Grant Program Award to X.W. (AUIGP 180271) and a generous laboratory start-up fund to X.W. from Auburn University College of Veterinary Medicine. This work is supported by the USDA National Institute of Food and Agriculture, Hatch project 1018100. X.X. and W.C. are supported by Auburn University Presidential Graduate Research Fellowship. Contributions of J.H.W. were supported by US NSF IOS 1456233 and the Nathaniel and Helen Wisch Professorship. We thank HudsonAlpha Genomic Services Lab for Illumina sequencing and Ting Li for help with DNA extractions.

## Supplemental Tables

**Table S1.**
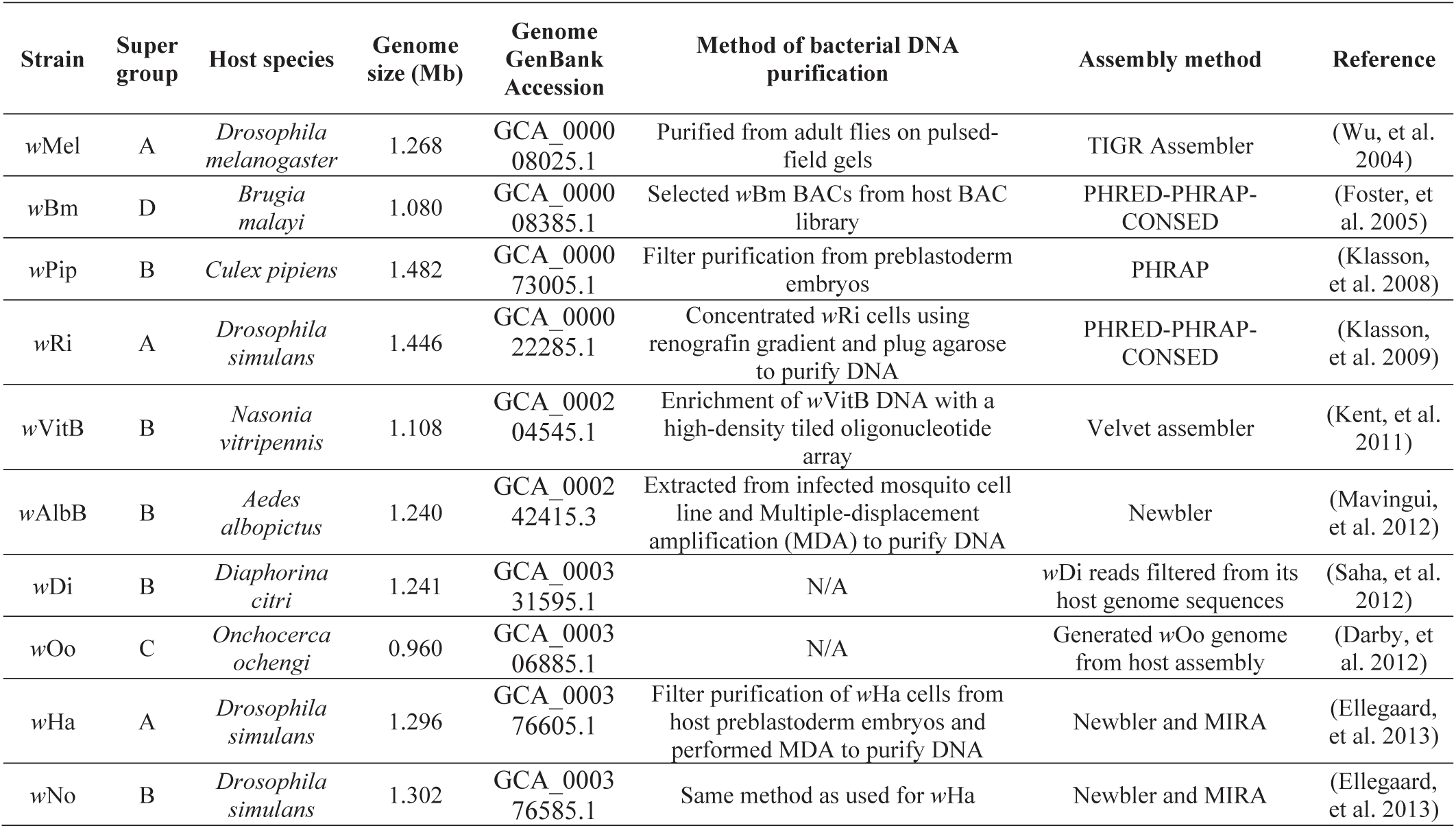
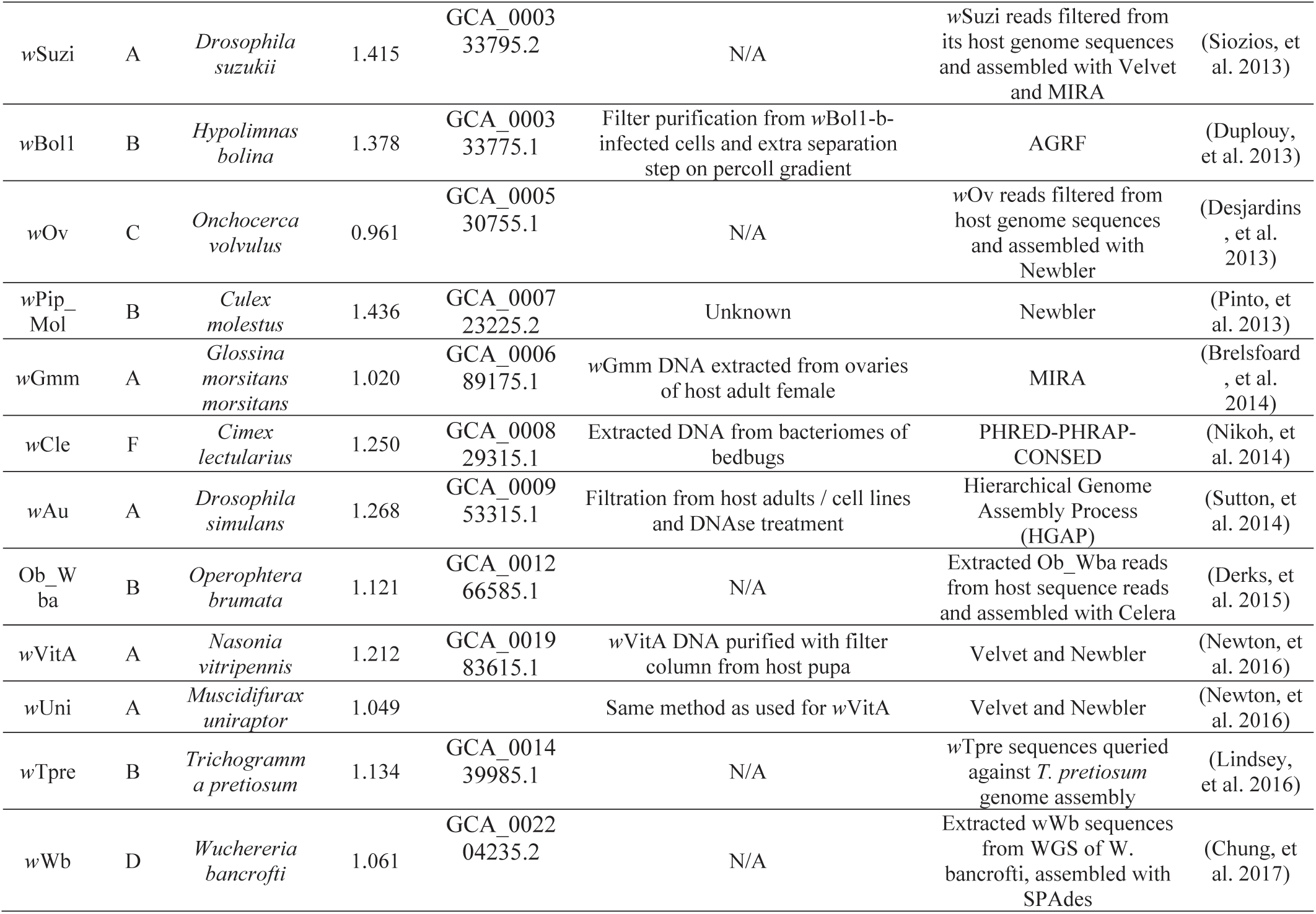
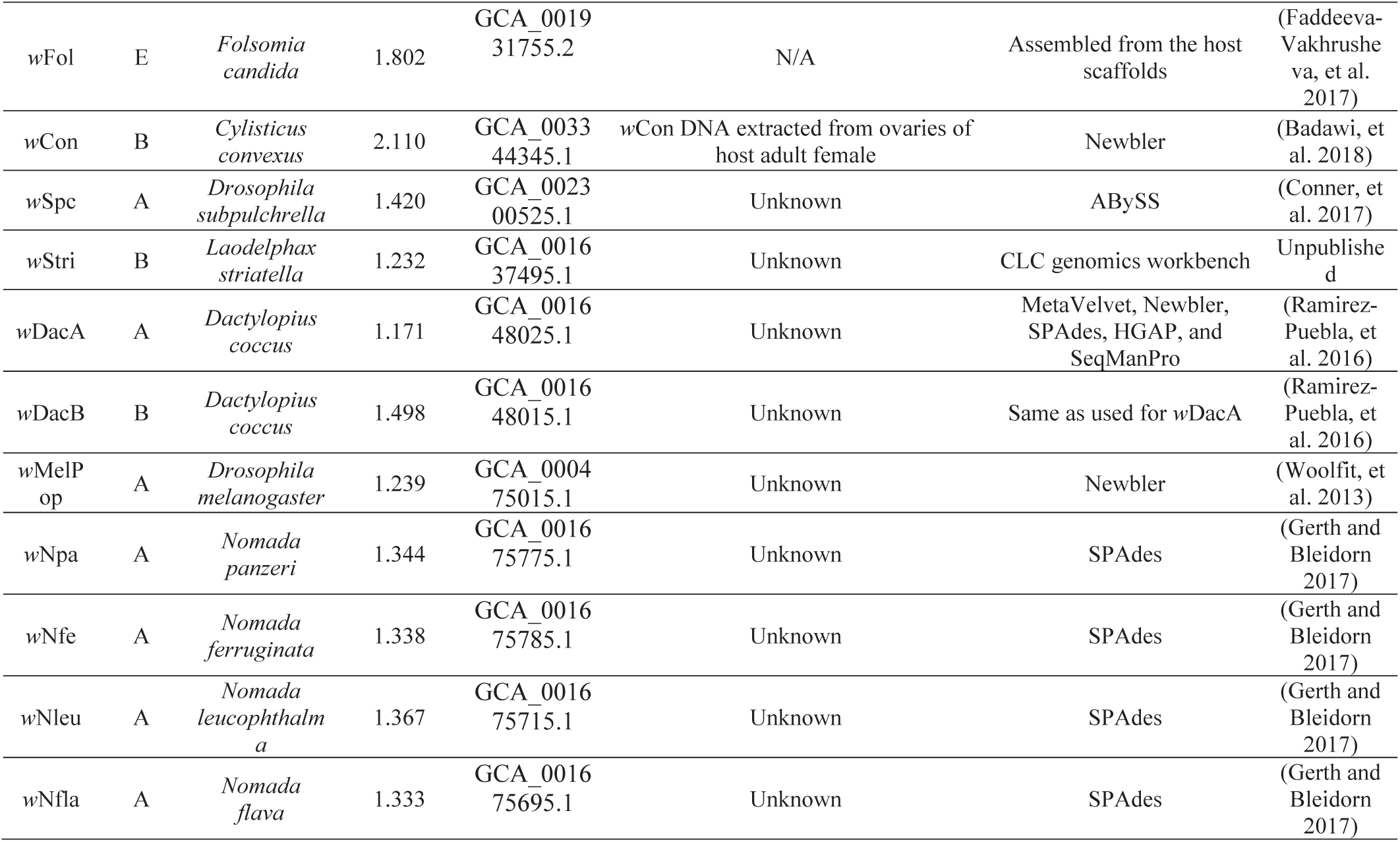
Summary of current sequenced *Wolbachia* whole genomes.

**Table S2.** Strain type of *w*One using informative SNPs in MLST genes.

| MLST gene | SNP position | A1 allele | A1 allele count | A2 allele | A2 allele count |
| --- | --- | --- | --- | --- | --- |
| gatB | 23 | G | 46 | A | 0 |
|  | 27 | A | 49 | G | 0 |
|  | 126 | T | 52 | C | 0 |
|  | 147 | G | 55 | A | 0 |
|  | 231 | A | 36 | G | 0 |
|  | 264 | T | 40 | C | 0 |
|  | 272 | T | 39 | C | 0 |
|  | 292 | A | 45 | C | 0 |
|  | 296 | G | 45 | A | 0 |
|  | 315 | C | 43 | T | 0 |
|  | 366 | G | 39 | T | 0 |
| coxA | 15 | C | 41 | A | 0 |
|  | 58 | T | 54 | C | 0 |
|  | 104 | T | 53 | C | 0 |
| hcpA | 126 | A | 48 | G | 0 |
|  | 132 | A | 43 | C | 0 |
|  | 148 | A | 42 | G | 0 |
|  | 150 | T | 44 | G | 0 |
|  | 179 | C | 51 | T | 0 |
|  | 180 | C | 52 | T | 0 |
|  | 192 | C | 51 | T | 0 |
|  | 201 | T | 54 | A | 0 |
|  | 225 | C | 55 | A | 0 |
|  | 234 | T | 57 | C | 0 |
|  | 237 | A | 57 | G | 0 |
|  | 246 | T | 57 | C | 0 |
|  | 249 | A | 61 | G | 0 |
| fbpA | 45 | C | 68 | T | 0 |
|  | 54 | G | 65 | A | 0 |
|  | 58 | A | 71 | G | 0 |
|  | 63 | G | 68 | A | 0 |
|  | 66 | T | 69 | C | 0 |
|  | 75 | A | 72 | G | 0 |
|  | 102 | A | 57 | G | 0 |
|  | 114 | C | 57 | A | 0 |
|  | 120 | G | 56 | A | 0 |
|  | 126 | T | 56 | C | 0 |
|  | 174 | A | 54 | G | 0 |
|  | 185 | A | 53 | G | 0 |
|  | 190 | G | 56 | A | 0 |
|  | 213 | G | 52 | A | 0 |
|  | 226 | T | 52 | C | 0 |
|  | 252 | A | 62 | G | 0 |
|  | 276 | G | 68 | A | 0 |
|  | 300 | A | 75 | G | 0 |
|  | 354 | A | 88 | G | 0 |
|  | 400 | C | 77 | T | 0 |
|  | 426 | A | 53 | G | 0 |

**Table S3.**
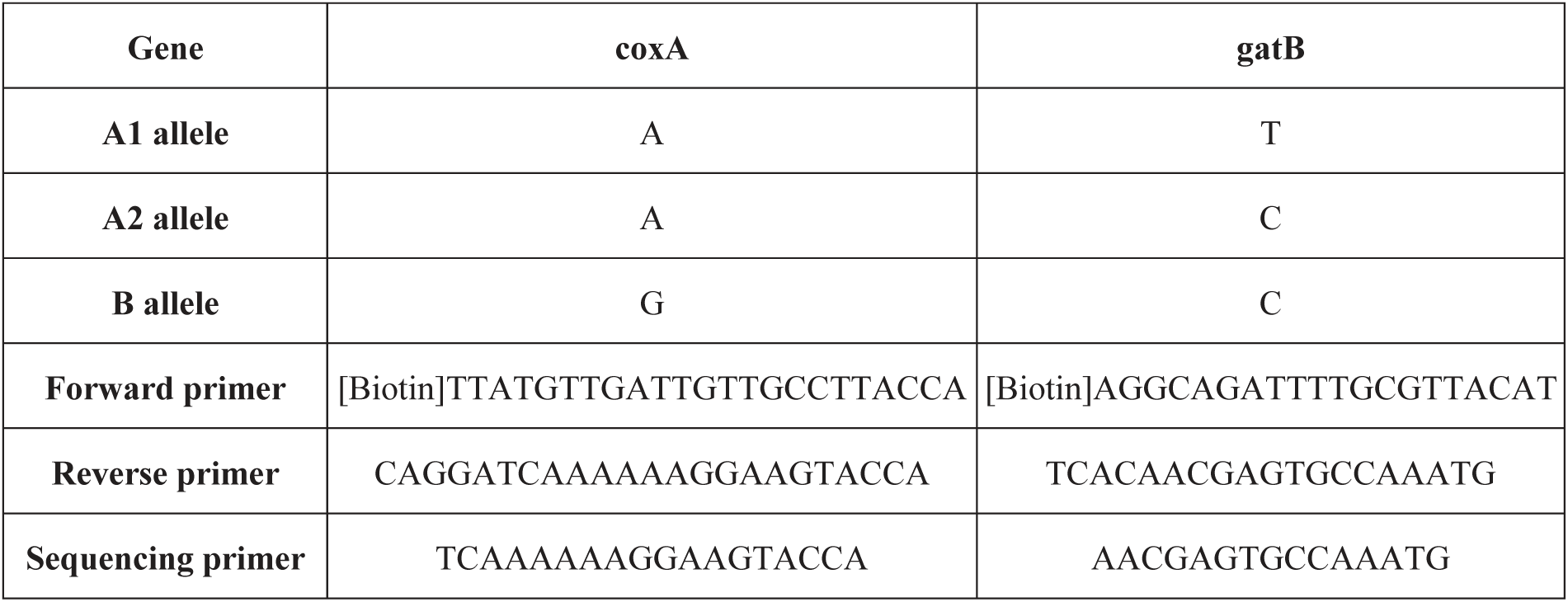
PCR and sequencing primers of coxA and gatB genes used in pyrosequencing for strain-typing of *Wolbachia* in *nasonia*.

**Table S4.** List of 27 genic segregating SNPs in *w*One genome.

| SNP ID | Scaffold | Position | Alleles | Coverage | Reference<br>allele count | Alternative<br>allele count | Gene ID | SNP effect |
| --- | --- | --- | --- | --- | --- | --- | --- | --- |
| wOneSNP03 | SCAFFOLD1 | 54805 | T/G | 84 | 57 | 27 | hypothetical protein WONE_00490 | nonsynonymous / moderate |
| wOneSNP07 | SCAFFOLD1 | 120377 | A/C | 88 | 64 | 24 | SecY | nonsynonymous / moderate |
| wOneSNP15 | SCAFFOLD1 | 208872 | A/G | 88 | 60 | 28 | hypothetical protein WONE_01750 | nonsynonymous / moderate |
| wOneSNP16 | SCAFFOLD1 | 208889 | G/T | 89 | 55 | 34 | hypothetical protein WONE_01750 | nonsynonymous / moderate |
| wOneSNP27 | SCAFFOLD2 | 197059 | A/G | 92 | 49 | 43 | hypothetical protein WONE_03520 | synonymous |
| wOneSNP30 | SCAFFOLD3 | 128108 | T/C | 80 | 48 | 32 | hypothetical protein WONE_04670 | synonymous |
| wOneSNP31 | SCAFFOLD3 | 128171 | C/A | 72 | 34 | 38 | hypothetical protein WONE_04670 | nonsynonymous / moderate |
| wOneSNP32 | SCAFFOLD3 | 128177 | C/T | 67 | 32 | 35 | hypothetical protein WONE_04670 | synonymous |
| wOneSNP33 | SCAFFOLD3 | 130774 | A/G | 96 | 75 | 21 | hypothetical protein WONE_04690 | synonymous |
| wOneSNP34 | SCAFFOLD3 | 156067 | G/C | 81 | 60 | 21 | cysS | nonsynonymous / moderate |
| wOneSNP39 | SCAFFOLD5 | 41230 | C/T | 92 | 69 | 23 | MFS transporter | nonsynonymous / moderate |
| wOneSNP44 | SCAFFOLD10 | 322 | T/A | 80 | 54 | 26 | baseplate assembly protein | synonymous |
| wOneSNP45 | SCAFFOLD10 | 331 | T/C | 82 | 54 | 28 | baseplate assembly protein | synonymous |
| wOneSNP46 | SCAFFOLD10 | 448 | C/T | 89 | 62 | 27 | baseplate assembly protein | synonymous |
| wOneSNP47 | SCAFFOLD10 | 611 | A/G | 91 | 52 | 39 | baseplate assembly protein | nonsynonymous / moderate |
| wOneSNP48 | SCAFFOLD10 | 636 | C/T | 94 | 61 | 33 | baseplate assembly protein | nonsynonymous / moderate |
| wOneSNP49 | SCAFFOLD10 | 642 | T/C | 94 | 62 | 32 | baseplate assembly protein | nonsynonymous / moderate |
| wOneSNP50 | SCAFFOLD10 | 647 | G/C | 90 | 62 | 28 | baseplate assembly protein | nonsynonymous / moderate |
| wOneSNP51 | SCAFFOLD10 | 877 | T/A | 81 | 45 | 36 | baseplate assembly protein | synonymous |
| wOneSNP52 | SCAFFOLD10 | 928 | C/A | 75 | 36 | 39 | baseplate assembly protein | synonymous |
| wOneSNP53 | SCAFFOLD10 | 931 | G/A | 75 | 36 | 39 | baseplate assembly protein | synonymous |
| wOneSNP54 | SCAFFOLD10 | 964 | G/A | 69 | 45 | 24 | baseplate assembly protein | synonymous |
| wOneSNP56 | SCAFFOLD18 | 11384 | T/C | 88 | 47 | 41 | hypothetical protein WONE_09680 | synonymous |
| wOneSNP57 | SCAFFOLD18 | 11387 | C/A | 93 | 45 | 48 | hypothetical protein WONE_09680 | synonymous |
| wOneSNP63 | SCAFFOLD30 | 5939 | C/T | 84 | 61 | 23 | hypothetical protein WONE_10710 | synonymous |
| wOneSNP65 | SCAFFOLD32 | 434 | A/G | 94 | 51 | 43 | hypothetical protein WONE_10780 | synonymous |
| wOneSNP66 | SCAFFOLD38 | 2021 | T/C | 66 | 42 | 24 | transposase | nonsynonymous / moderate |

## Supplemental Figures

**Figure S1.**
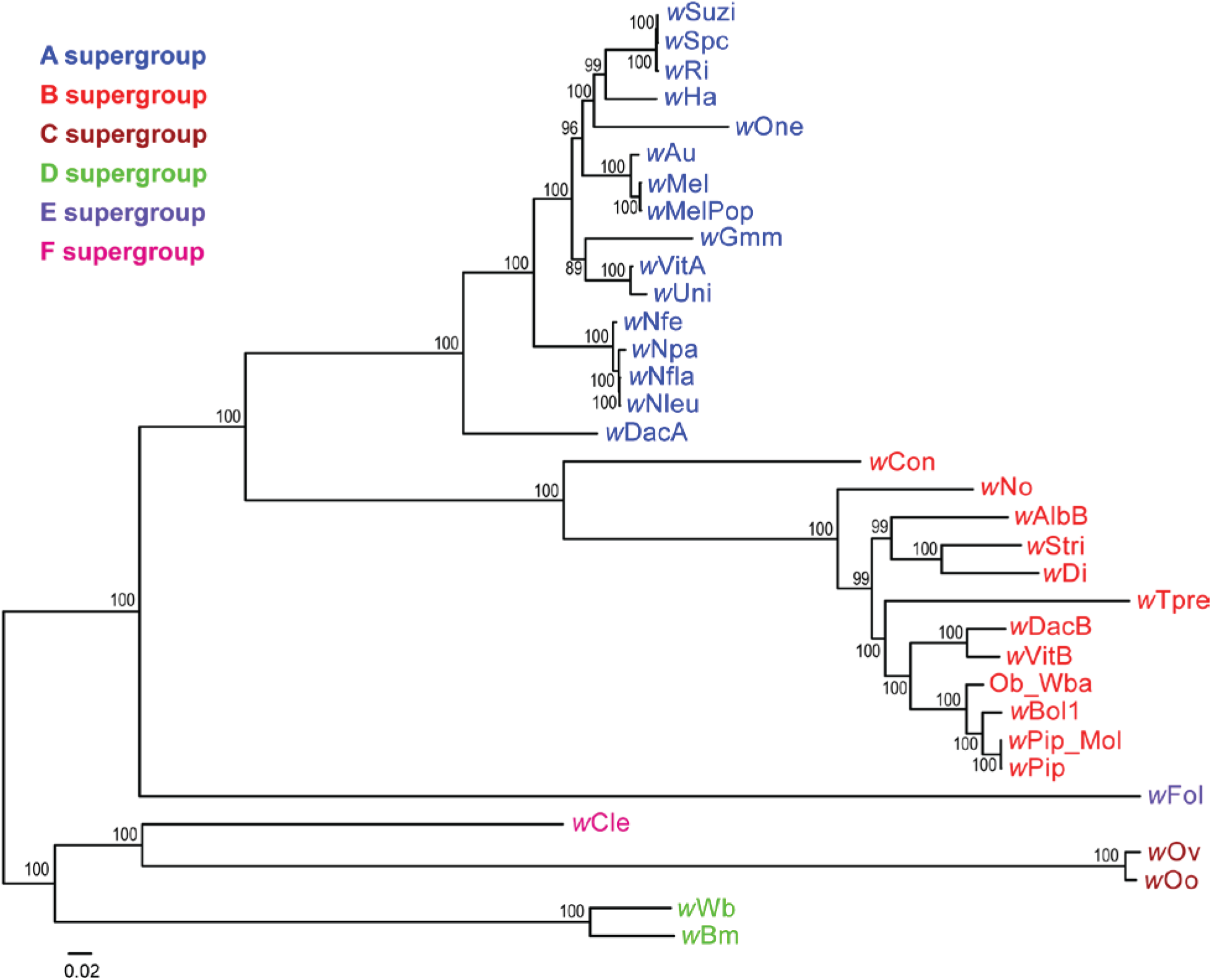
Phylogenetic analysis of 34 *Wolbachia* genomes using Maximum Likelihood method from a concatenated protein alignment of 211 single-copy orthologous genes.

**Figure S2.**
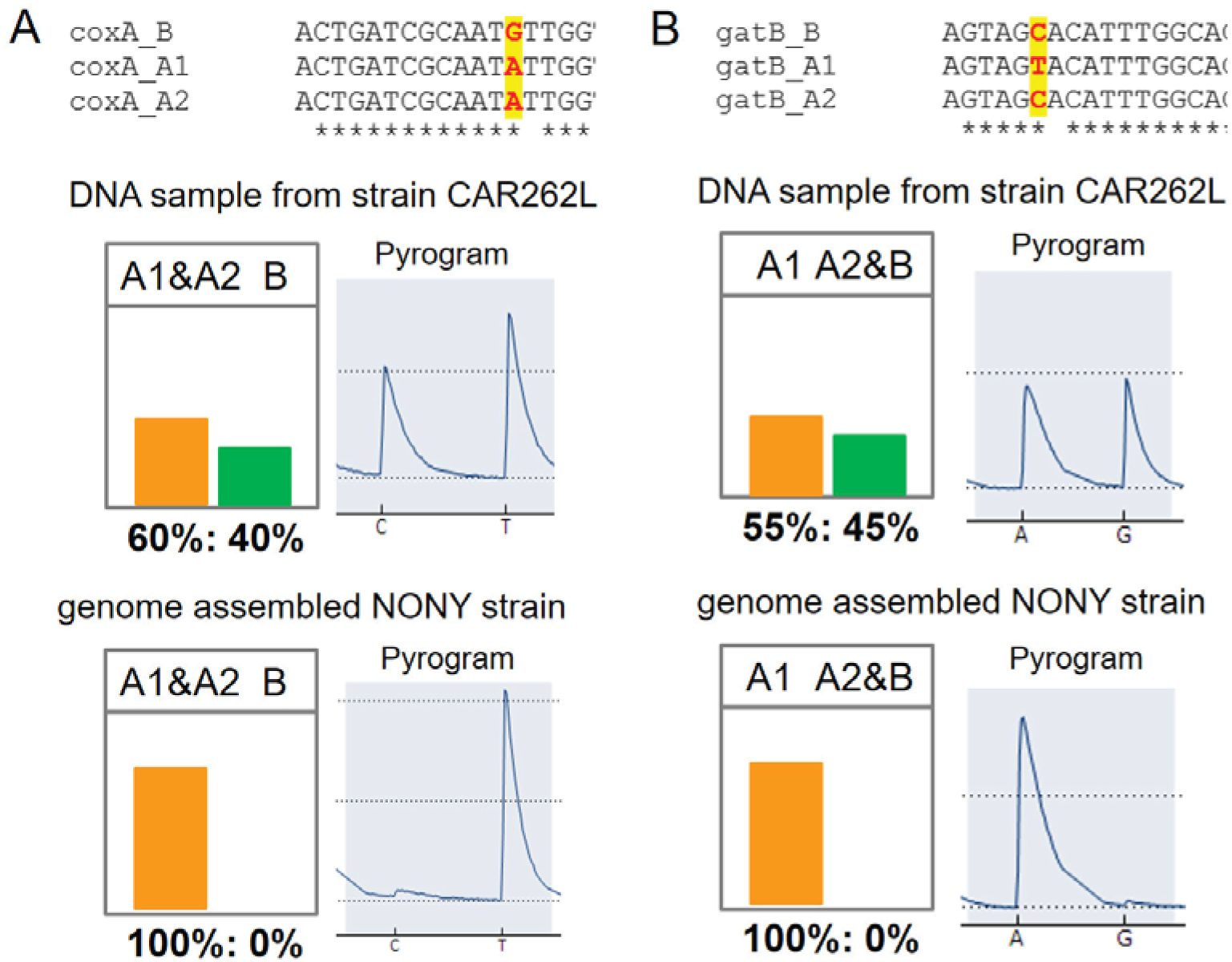
Allele-specific pyrosequencing for *Wolbachia* strain typing in genome sequenced *N. oneida* and the wild strain CAR262L.

**Figure S3.**
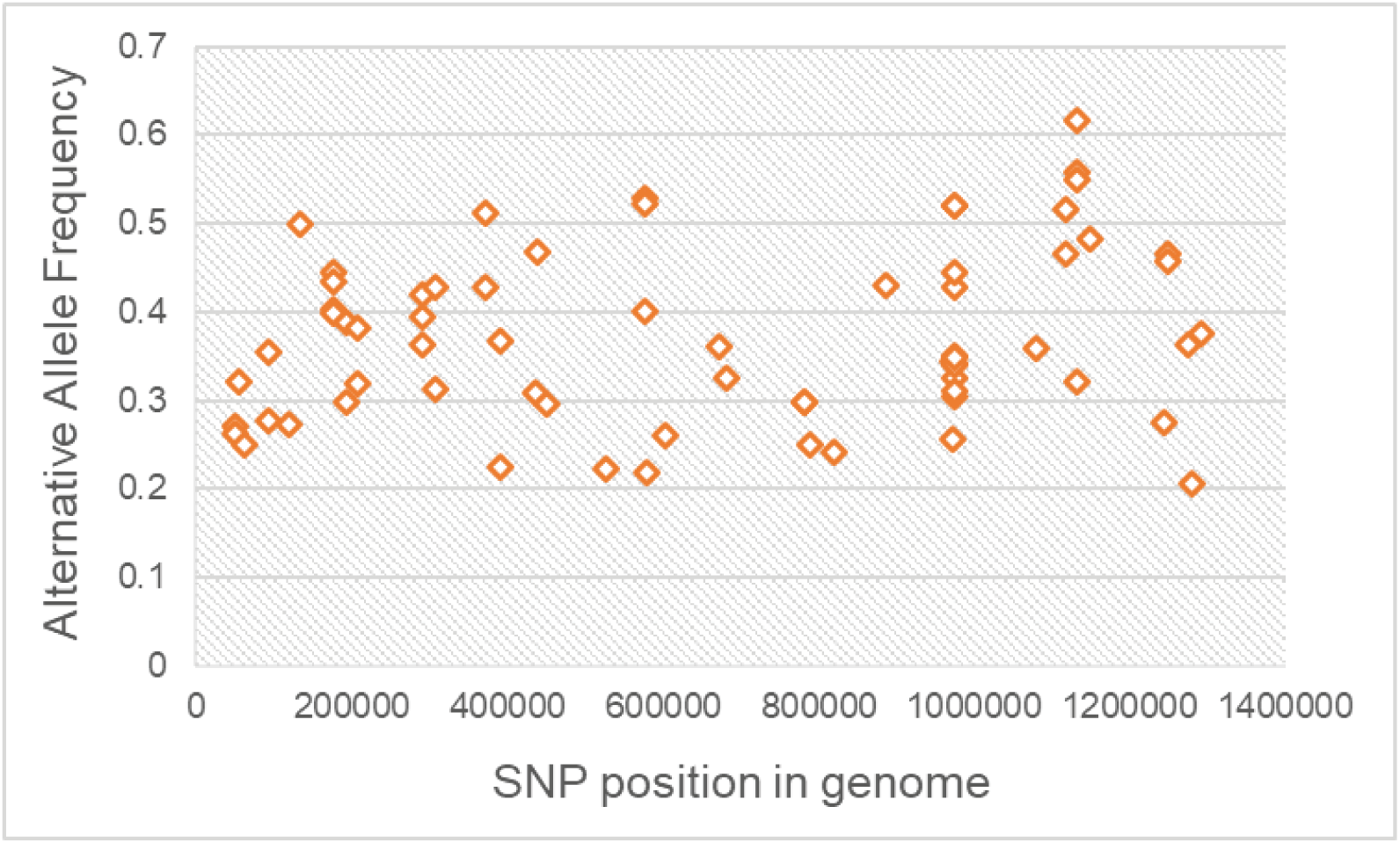
Alternative allele frequency of SNPs identified in *w*One genome.

**Figure S4.**
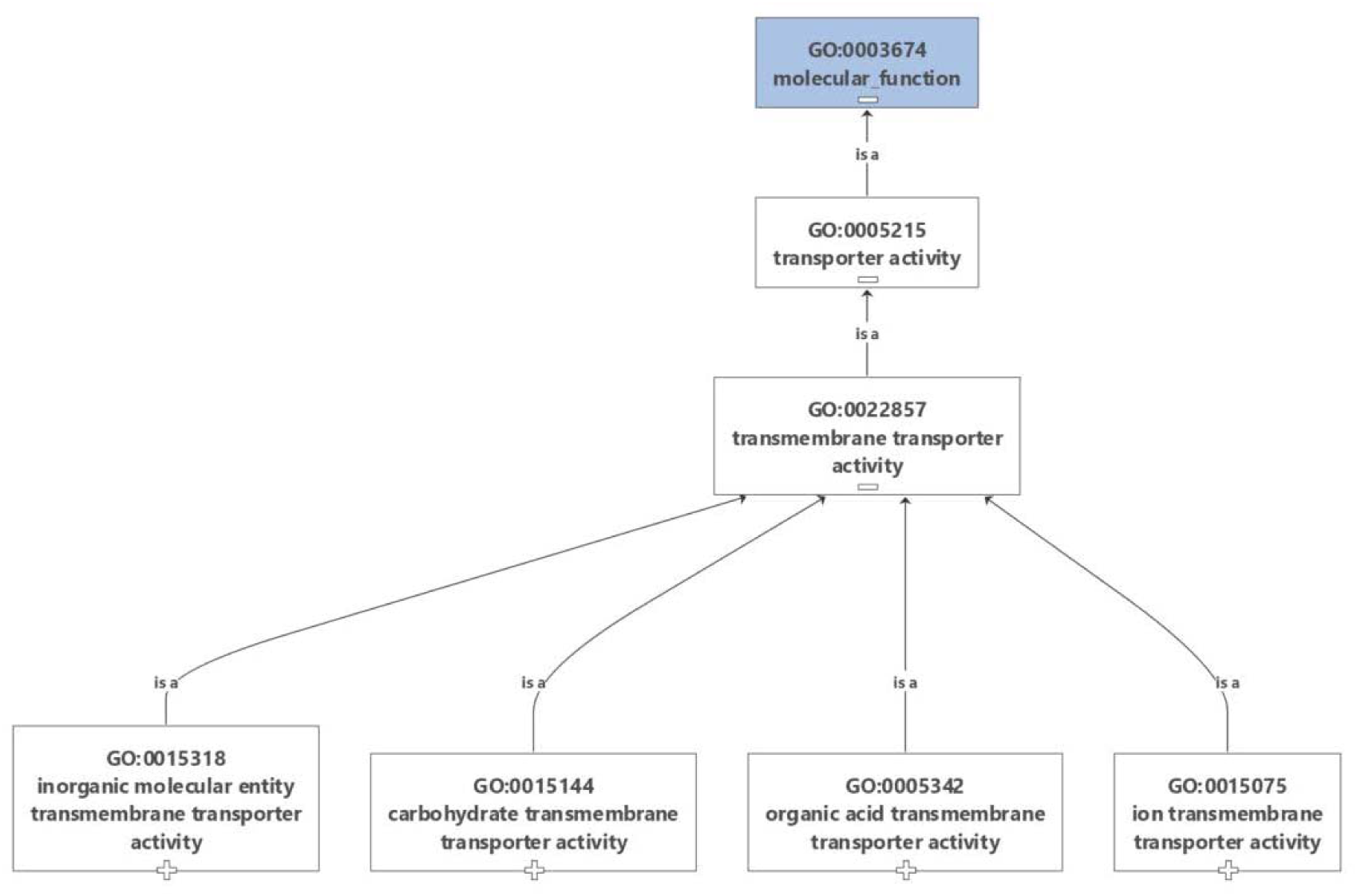
Enriched GO terms of genes with identified SNPs in the *w*One genome.

**Figure S5.**
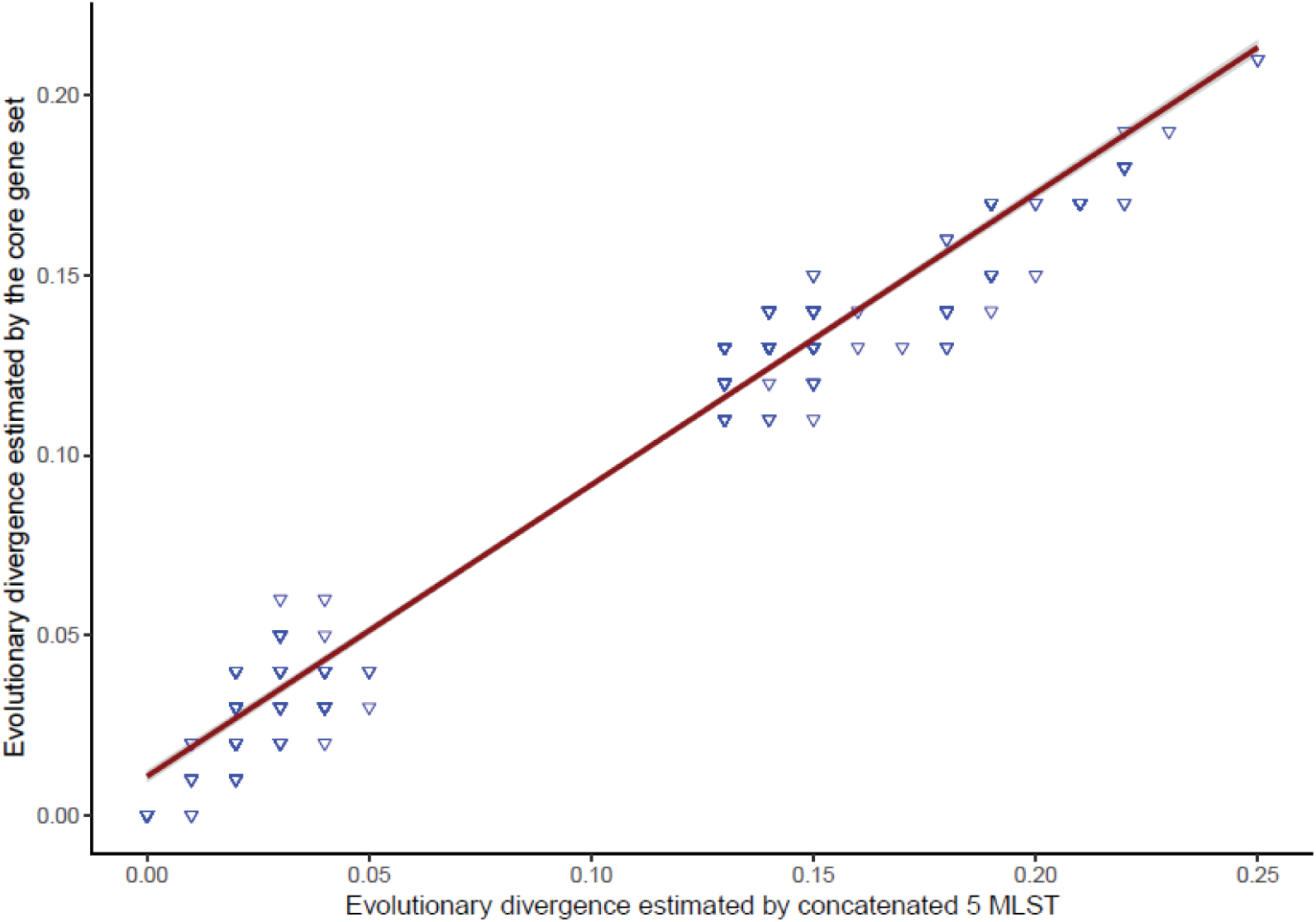
Correlation of evolutionary divergence estimated by core gene set and concatenated five MLST genes.

